# Modeling Psychophysical Data in R: A Comparative Study of Four Model Frameworks

**DOI:** 10.1101/2025.11.17.688833

**Authors:** Priscilla Balestrucci, Maura Mezzetti, Barbara La Scaleia, Alessandro Moscatelli

## Abstract

Inferential models in psychophysics are essential for quantifying the relation between physical properties of the stimulus and their perceptual representations. The psychometric function is typically used to model the responses of individual participants in forced-choice experiments. The accuracy and the noise of the response can be estimated from the Point of Subjective Equality (PSE) and the Just Noticeable Difference (JND) of the function, respectively. Traditionally, a two-level approach is used to model the behavior of a group of participants, where psychometric functions are first fitted to individual participant data, followed by hypothesis testing across participants on parameters of interest. Recent studies have introduced alternative approaches based on hierarchical models, such as Generalized Linear Mixed Models (GLMM) and Models within the Bayesian Hierarchical Framework (BHF), to analyze in a single framework the responses from multiple participants. These two approaches can be effectively implemented in R, thanks to its flexibility and robust statistical capabilities. Here, we provide a tutorial on how to model and analyze data from psychophysical experiments in R, using both two-level and hierarchical frameworks. Our goal is to provide researchers with a practical guide for building a complete and reproducible analysis pipeline, using core R functionalities together with custom packages, and facilitate rigorous and efficient data analysis in psychophysics.

## Introduction

Psychophysics investigates the quantitative relationship between some physical properties of the environment and their perceptual representation provided by the senses. For example, the perceived brightness of a visual stimulus (Treisman, 1964), the speed of an object sliding across the skin (Dallmann et al., 2015; Ryan et al., 2022), or the pitch of a sound (Senna et al., 2015), are properties of a stimulus that can be investigated with a psychophysical experiment. Psychophysical studies provide insights into fundamental sensory processes (Ernst & Banks, 2002; Moscatelli et al., 2019), offer quantitative assessments of sensory abilities under physiological and pathological conditions (Picconi et al., 2022; Fitze et al., 2024), and inform the design and evaluation of technological devices (Travis, 1990; Fani et al., 2017; Blau et al., 2024). Yes-no Tasks, Forced-Choice, 2-Alternative-Forced Choice (2AFC), or n-Alternative Forced Choice (n-AFC) are examples of psychophysical tasks that researchers use to infer the internal processes underlying the perceptual representation of a stimulus (Pelli & Bart, 1991; Wichmann et al., 2018).

In this article we will compare four model frameworks to analyze psychophysical data. Models will be evaluated on two datasets from real and simulated experiments. Usually, binary responses of forced choice experiments are modeled at the participant level using the psychometric function (Wichmann & Hill, 2001a,b), which has a sigmoidal shape and relates the intensity of a test stimulus to the probability of the response. The parameters of the psychometric function summarize the observer’s performance and the characteristics of the underlying sensory mechanisms. Two key parameters are the Point of Subjective Equality (PSE) and the Just Noticeable Difference (JND). The PSE is the stimulus value at which an observer is equally likely to judge a test stimulus as greater or lesser than the reference (typically at 50% response probability on the psychometric function). It represents the point where the test stimulus is perceived as equal to the reference, even if their physical values are not identical. If the value of the PSE differs from the value of the reference stimulus, it may indicate a perceptual bias. The JND is a measure of sensitivity—the smallest change in stimulus intensity that an observer can reliably detect. On a psychometric function, it is typically defined as half the difference between the stimulus values corresponding to response probabilities of 25% and 75%. This captures the steepness of the curve: a smaller JND means the observer can discriminate finer differences. Other parameters that are sometimes introduced in the model are the guess (*γ*) and lapse rate (*λ*), which define the lower and upper asymptote of the psychometric curve, respectively (Wichmann & Hill, 2001a). The guess rate represents the probability of a correct response when responding at chance, and its value is largely determined by the experimental procedure used (*γ* = 0 in yes-no tasks, *γ* = 1*/n* in *n*-AFC paradigms). The lapse rate represents the probability of an incorrect response regardless of stimulus intensity, e.g. due to a drop in vigilance or an error in signaling the intended response, and does not depend on the experimental design. Although these two parameters are not directly used in characterizing the sensory mechanisms involved, their wrong assumption or estimation can lead to a biased estimation of the other characteristics of the psychometric function (Wichmann & Hill, 2001a; Yssaad-Fesselier & Knoblauch, 2006; Prins, 2012; Klein, 2001; Zchaluk & Foster, 2009).

Different statistical methods have been proposed for fitting and analyzing psychophysical data. A widely used approach for modeling psychometric functions is the Generalized Linear Model (GLM, Agresti (2002)), which enables estimating the PSE and the JND at the individual participant level (Knoblauch & Maloney, 2012). By using the GLM approach, however, it is not possible to estimate or fix the asymptotes of the function, preventing the inclusion of lapse and guess rates in the model. Other approaches, such as maximum likelihood methods based on generalized non-linear models (GNM, Yssaad-Fesselier & Knoblauch (2006)) or Bayesian inference methods (Kuss et al., 2005), allow for the estimation of additional parameters of the psychometric function at the level of the individual participants. While modeling individual responses is essential, generalizing findings to the population level often requires a second layer of analysis. Therefore, after fitting psychometric functions for each individual observer, it is typically necessary to aggregate processed data (e.g., PSE or JND estimates) across participants and perform inferential statistics, for example using ANOVA or t-tests.

In recent years, alternative approaches have been introduced to estimate and analyze responses from multiple participants at the population level. Notably, Generalized Linear Mixed Models (GLMMs, Moscatelli et al. (2012)) and Bayesian Hierarchical Models (Mezzetti et al., 2023; Prins, 2024; Fitze et al., 2024; Courtin et al., 2025) allow for simultaneous estimation of both individual- and group-level parameters. These methods offer advantages over traditional two-level approaches, such as higher statistical power (Moscatelli et al., 2012) and, in the case of Bayesian models, the possibility to incorporate prior knowledge from existing literature and past studies to reduce model uncertainty (Mezzetti et al., 2023).

The analysis of data from psychophysical experiments can be implemented in various programming languages and environments, with R, Python, Julia, and MATLAB being the most widely used. In addition to general-purpose statistical computing libraries, several tools have been specifically designed for psychophysical analysis. Notable examples include *MPDiR* (Knoblauch & Maloney, 2012), *psyphy* (Knoblauch, 2023), *and quickpsy* (Linares et al., 2016) *in R; psignifit* (Wichmann & Hill, 2001a,b) *in MATLAB and Python; Palamedes* (Prins & Kingdom, 2018) *in MATLAB; and PsychoPy* (Peirce, 2007) in Python. Probabilistic programming languages such as Stan, JAGS, and PyMC, with interfaces for R, MATLAB and Python provide flexible frameworks for specifying and fitting Bayesian models commonly used in psychophysical data analysis.

This tutorial aims to guide researchers in selecting and implementing appropriate statistical models for analyzing psychophysical data, with an emphasis on R-based tools. We developed our own R package *MixedPsy* (Moscatelli & Balestrucci, 2021) providing tools for the analysis of psychophysical data, which is available for download on CRAN and GitHub. Throughout this paper, we will use functions from *MixedPsy* as well as from base R and other packages, to demonstrate different pipelines for the analysis of psychophysical datasets.

We provide an overview of several models that can be used to estimate psychophysical parameters. At the individual participant level, we introduce the Generalized Linear Model (GLM) and the Generalized Nonlinear Model (GNM). Among hierarchical molders, we cover the Generalized Linear Mixed Model (GLMM), and Bayesian Hierarchical Models. We consider two possible implementations of Bayesian Hierarchical Models: one model type with the asymptotes fixed to zero, labeled Bayesian Hierarchical GLM (BH-GLM), and one with specific parameters to model asymptotes different from zero, labeled Bayesian Hierarchical GNM (BH-GNM). For the sake of clarity, a list of the acronyms most frequently used throughout this paper is provided in Table 1. We illustrate each model with two example datasets from forced-choice discrimination tasks, where participants are requested to compare the magnitude of a test to a reference stimulus. The first is a simulated dataset with known parameters, allowing for direct comparison between estimated and true values. The second is an empirical dataset from an experiment reported by Dallmann et al. (2015), which has also served as a benchmark in Mezzetti et al. (2023). The equations of these model frameworks are briefly described in the Methods section. In the Results section, we described the R and Stan (for BH-GLM and BH-GNM) syntax to fit the five models to the data. Finally, the estimates of the models for each dataset are evaluated in the Model Diagnostics and Statistical Inference paragraphs.

**Table 1.** Acronyms frequently used in the paper. The Bayesian Hierarchical Framework included the two model types BH-GLM and BH-GNM.

| Acronym | Definition |
| --- | --- |
| PSE | Point of Subjective Equivalence |
| JND | Just Noticeable Difference |
| GLM | Generalized Linear Model |
| GNM | Generalized Non-Linear Model |
| GLMM | Generalized Linear Mixed Model |
| BHF | Bayesian Hierarchical Framework |
| BH-GLM | Bayesian Hierarchical GLM |
| BH-GNM | Bayesian Hierarchical GNM |

## Methods

### Datasets

#### Example 1: A simulated dataset

The simulated dataset `simul_data` has been generated using the *MixedPsy* function `PsySimulate()`. Five example rows of the dataset are shown in Table 2. It mimics an experiment of time perception based on the method of constant stimuli, where ten observers (identified in the `Subject` column) compare two stimuli of different duration, reference and test stimulus, and judge which one last longer. Despite the use of a length-based label, the dataset is structured to emulate a generic forced-choice discrimination paradigm. The independent variable `X` represents duration of the comparison stimulus with nine equally spaced levels ranging from 40 to 120 arbitrary units (e.g., milliseconds). The binomial response is recorded in the `Longer` and `Total` columns, which indicate how many times the test stimulus at a given duration was judged as longer than the reference stimulus, and the total number of trials in which that duration value was presented, respectively.

**Table 2.** Example of observations from the simulated dataset.

| Subject | X | Intercept | Slope | Gamma | Lambda | Longer | Total |
| --- | --- | --- | --- | --- | --- | --- | --- |
| 1 | 40.00 | -7.87 | 0.09 | 0.00 | 0.01 | 0 | 160 |
| 2 | 100.00 | -4.59 | 0.09 | 0.02 | 0.02 | 157 | 160 |
| 4 | 60.00 | -6.29 | 0.09 | 0.01 | 0.02 | 28 | 160 |
| 5 | 70.00 | -8.06 | 0.09 | 0.03 | 0.01 | 14 | 160 |
| 6 | 120.00 | -5.10 | 0.09 | 0.01 | 0.01 | 159 | 160 |

The dataset can be generated using the *MixedPsy* function `PsySimulate()` as follows:

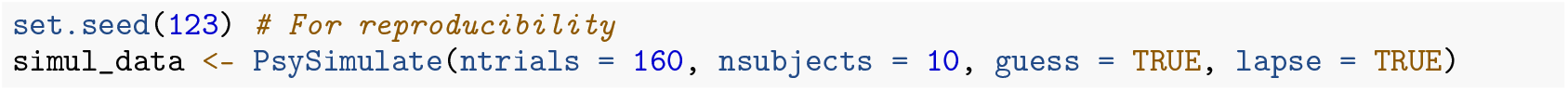

Using the `PsySimulate()` function, it is possible to simulate a variety of responses by modifying parameters such as the fixed and random effects matrices, the number of subjects, the range of the independent variable, and the simulation of lapse and guess rates. Ten simulated participants were sampled from multivariate normal distribution *𝒩* (*θ*, Σ) with *θ*_*x*_ = *−*7 (Intercept) and *θ*_*y*_ = 0.0875 (Slope)—corresponding to *PSE* = 80 amd *JND* = 7.7. For the specific seed indicated in the R code, the sample mean was equal to 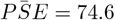 and the 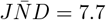, hereafter referred to as the true values of PSE and JND. The lapse and guess rates vary for each observer are drawn from a uniform distribution between 0 and 0.05. Example of simulated observation and parameters are shown in Table 2.

### Example 2: The role of vibration in tactile speed perception

The dataset named `vibro_exp3` is part of a study on tactile discrimination of surface motion (Dallmann et al., 2015). In the experiment, nine participants judged the speed of a moving surface sensed by touch while exposed to different vibratory noise conditions. In each trial, participants evaluated whether motion of a test stimulus was faster or slower than a reference. The test stimulus speed ranged between 1.0 and 16.0 cm/s across seven equally spaced values, and the masking vibrations were either absent (0 Hz, control condition) or had a frequency of 32 Hz (masking condition). Each combination of speed and vibration was tested 40 times in randomized order, resulting in a total of 560 trials per participant.

The dataset is organized into 5 columns, each corresponding to one of the variables of interest, namely: the two independent variables, test stimulus speed (`speed`) and masking vibration frequency (`vibration`); the response variables indicating the number of trials in which the test stimulus was perceived as faster or slower than the reference (`faster` and `slower`, respectively); and the observer ID (`subject`). Table 3 shows five example rows of the dataset.

**Table 3.** Example of observation from the experimental dataset.

| speed | vibration | faster | slower | subject |
| --- | --- | --- | --- | --- |
| 3.50 | 0 | 2 | 38 | MA |
| 6.00 | 32 | 10 | 30 | MA |
| 11.00 | 0 | 26 | 14 | MI |
| 13.50 | 32 | 33 | 7 | NN |
| 8.50 | 32 | 18 | 22 | NN |

The dataset is included in the *MixedPsy* package.

## Models

### Generalized Linear Models

GLMs are commonly used to model the relationship between the stimulus intensity, *x*_*ij*_, for participant *i* and trial *j* and the probability of a binary response, *P* (*Y*_*ij*_ = 1 | *x*_*ij*_). The binary response *Y*_*ij*_ *∈* [0, 1] is for example “comparison faster” or “comparison slower” in the Forced-Choice discrimination task described in Dallmann et al. (2015), “Yes” or “No” in a detection task, etc. Let *π*_*ij*_ = *P* (*Y*_*ij*_ = 1 | *x*_*ij*_) be the probability of responding 1 to stimulus *x*_*ij*_, and the Bernoulli distribution of *Y*_*ij*_ defined as:

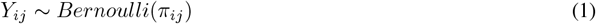

The model equation is the following:

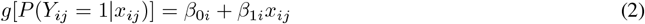

The link function *g*(*·*) allows to establish a linear relationship between the probability of a response and the continuous predictor, which is fully described by two parameters of intercept *β*_0*i*_ and slope *β*_1*i*_. Typical link functions are the Probit and Logit functions, which map the probability of the response through the inverse cumulative normal distribution function (labeled as Φ^*−*1^(*·*) below) and inverse cumulative logistic distribution functions, respectively (Agresti, 2002; Klein, 2001).

Parameters relative to observer *i*, PSE_i_ and JND_i_, are computed from intercept and slope in Eq.(2):

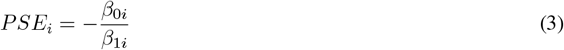

and

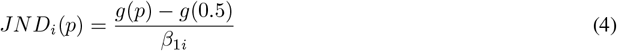

The numerator in Eq.(4) represents the difference in the link function evaluated at *p* (i.e. the p-quantile of the distribution) and at the point of subjective equality which corresponds to *p* = 0.5. In the case of a probit link function, *g*(*p*) is the quantile function of the standard normal distribution, so that *g*(0.5) = 0. A common choice is *p* = 0.75, which yields 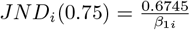. Alternatively, the JND can be calculated at the 84th percentile (*p* = 0.84), which corresponds approximately to one standard deviation above the mean of the normal distribution, 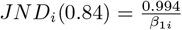

It is possible to rewrite Eq (2) with respect to the probability *P* (*Y*_*ij*_ = 1| *x*_*ij*_), either as a function of (*β*_0*i*_, *β*_1*i*_) or as a function of the location and scale parameters (*µ*_*i*_, *σ*_*i*_), which exactly correspond to *PSE*_*i*_ and 1*/β*_1*i*_, respectively:

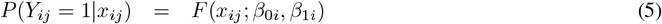

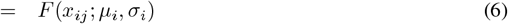

where *F* (*·*) = *g*^*−*1^(*·*) represents the inverse link function. The function *F* (*·*) correspond to the Cumulative Distribution Function (CDF) of a normal distribution in the probit model:

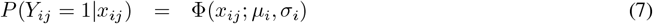

where *F* (*·*) = Φ (*·*) is the CDF of a normal distribution with mean *µ*_*i*_ = *PSE*_*i*_ and standard deviation *σ*_*i*_ = 1*/β*_1*i*_. Likewise, in logit models it corresponds to the CDF of a logistic distribution.

A key limitation of GLM is their assumption of fixed lower and upper asymptotes to 0 and 1, respectively—i.e., the range of a CDF. This makes them unsuitable for modeling data where guess and lapse rates must be accounted for—that is, when responses deviate from (0, 1) at very low and very high stimulus intensity, respectively. Another limitation lies in the assumption of independence across subjects, resulting in separate parameter estimates for each individual and the absence of population-level estimates.

### Generalized Non-Linear Models

To include guess and lapse rates, a more general form of the psychometric function in Eq. (5) can be written as in Wichmann & Hill (2001a):

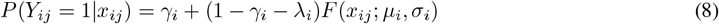

The parameter *γ*_*i*_ represents the lower asymptote (i.e., the guess rate). In n-AFC this is often fixed to the chance level 1*/n*. The parameter *λ*_*i*_ refers to the distance from the upper asymptote, accounting for stimulus-independent mistakes at high stimulus levels. The estimation of the model in Eq. (8) is more complicated than that of (2) due to nonlinearity; nevertheless, the model in Eq. (8) can be fitted in R using the GNM framework, which extends GLMs by allowing nonlinear relationships between predictors and responses (Yssaad-Fesselier & Knoblauch, 2006).

The PSE is equal to:

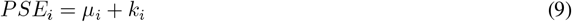

where *k*_*i*_(*γ*_*i*_, *λ*_*i*_) = 0 if *γ*_*i*_ = *λ*_*i*_ (see Supplementary Information for details). In forced choice experiments typically *γ*_*i*_ *≈ λ*_*i*_ and the value of *k*_*i*_ is negligible. Therefore, we approximated the PSE to Eq. (3) in both, GNM and BH-GNM. Similarly, the JND can be approximated to Eq. (4) for small values of *γ*_*i*_ and *λ*_*i*_. Analytical expressions for the PSE and JND in GNM and BH-GNM including *γ*_*i*_ and *λ*_*i*_ are provided in the Supplementary Information.

### Generalized Linear Mixed Models

In psychophysics experiments, data are naturally divided in clusters, since multiple responses are collected from each observer. Intuitively, repeated responses from a single participant are more correlated with each other than with those from other participants. GLMMs extend GLMs by accounting for clustered data structures (Agresti, 2002; Moscatelli et al., 2012). They do so by simultaneously modeling the fixed effects associated with the experimental variables and the random effects that capture variability across clusters (i.e., the differences between participants). The GLMM extends the psychometric function in Eq. (2) to account for multiple participant:

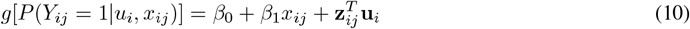

Model in Eq. (10) takes into account a single fixed effect predictor, *x*_*ij*_, and the random effect vector **u**_*i*_. In its general form, the model equation is formalized as follows:

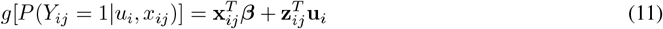

where **x**_*ij*_ is a vector of predictor variables for participant *i* and observation *j*; ***β*** is the vector of fixed effects, representing the parameters modeling the effects of the predictor variables at the population level; **z**_*ij*_ is the vector of explanatory variables associated with random effects—typically a subset of **x**_*ij*_; and **u**_*i*_ is the vector of random effects, which describe the deviations from the fixed-effects parameters for participant *i* and model the individual differences among participants in the population sample. In the model, random-effect parameters are assumed to be random variables with multivariate normal distribution *N* (**0, Σ**). The covariance matrix **Σ** describes the variance and covariance structure of the random effects and depends on the specifications of the model.

Using the GLMM in Eq. (11), it is possible to include different experimental conditions in the model, and possible interactions. For example, it is possible to evaluate, within a single model, how the probability of a binary response varies with changes in stimulus intensity across different experimental conditions. These conditions can be represented as levels of a categorical variable, and the model can be formalized as:

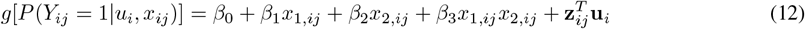

where *x*_1_ represents stimulus intensity, and *x*_2_ is a dummy-coded variable encoding for different conditions of a categorical predictor (e.g., *x*_2,*ij*_ = 0 for condition 1 and *x*_2,*ij*_ = 1 for condition 2). In this coding scheme, *β*_0_ and *β*_1_ represent the intercept and slope, respectively, associated with condition 1 (i.e., the reference condition), while *β*_2_ and *β*_3_ describe the change in intercept and slope for condition 2 relative to condition 1.

From the fixed-effect parameters of the model in Eq. (10), represented by the intercept *β*_0_ and the slope *β*_1_, PSE and JND can be derived using the same relationship described in Eq. (3) and (4). In this case, however, the resulting estimates refer to the performance at the population level, rather than of individual participants. Estimates of PSE and JND for the different conditions in (12) can be derived considering the dummy-coded structure of the model coefficients as follows:

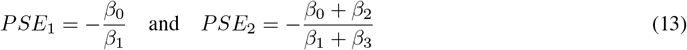

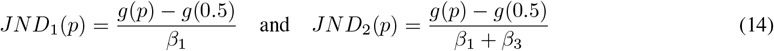

The specification of the random-effects structure is a crucial aspect of model building, as it influences both model fit and the precision of the fixed-effect estimates. The inclusion of random effects allows accounting for individual heterogeneity, thereby providing a better evaluation of the precision of the fixed-effect estimates, as well as yielding population-level estimates.

In principle, a GLMM can incorporate any combination of fixed and random effects. However, it is generally advisable to avoid overly complex models to improve interpretability and reduce the risk of overfitting. Overfitting occurs when a model captures not only the signal, but also the random noise in the data, which can impair its ability to generalize to new samples from the same population James et al. (2013).

### Bayesian Hierarchical Models

In accordance with the work in Mezzetti et al. (2023), we first propose a two-stage bayesian hierarchical model where the first level coincides with Eq. (2) (BH-GLM). We define the Bernoulli distribution for the response variable *Y* as in Eq. (1).

For each observer, *i*, the response probability, *P* (*Y*_*ij*_ = 1| *x*_*ij*_) = *π*_*ij*_, is linked with the explanatory variable by a probit equation as in Eq. (2).

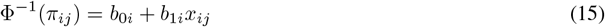

The second level of the hierarchical model integrates both individual and population-level components. It effectively assesses overall effects across subjects by incorporating individual-specific variations. The parameters (*β*_0_, *β*_1_) define the population model, obtained by combining the subject-specific parameters while accounting for their associated uncertainties, incorporated in the second level of the model:

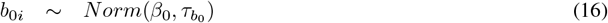

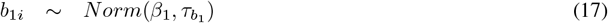

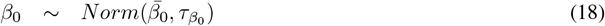

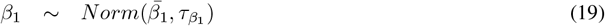

In line with the recommendations of Gelman (2006); Gelman et al. (2008), who advocate the use of this prior to mitigate overfitting and underfitting, a Cauchy distribution with location parameter 0 and scale parameter 2.5 is employed as a robust and weakly informative choice for the hyperprior distributions of 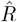. The Cauchy distribution has heavy tails, meaning it does not overly penalize large values of the standard deviation. This is useful when one does not want to impose a too restrictive prior, allowing the data to drive the estimation. In general, taking a value of the scale hyperparameter equal to 2.5, unlike a uniform prior (which can lead to inference issues), favors small values but does not exclude large ones.

Eq. (15) can be redefined directly through parameters *pse*_*i*_ = *µ*_*i*_ and *σ*_*i*_ as:

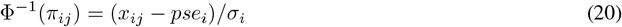

Similarly, the individual parameters are linked with the with the population-level parameters:

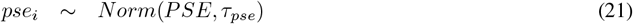

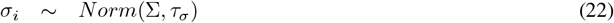

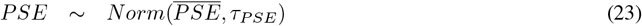

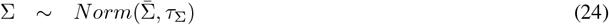

Ideally, a prior distribution represents the degree of confidence scientists have in the true model parameters. When empirical data are available, new information can be systematically integrated through statistical models using Bayesian learning. This process begins by capturing existing expert knowledge and associated uncertainty. In general, prior distributions for the model parameters can be specified as noninformative (as the ones used in Example 1 and in Example 2), subjective prior reflecting an informed estimation of a parameter’s value before any data are collected, or informed by historical data such as in the power prior approach. Following Ibrahim & Chen (2000), the power prior framework incorporates information from a historical dataset *D*_0_ = (*n*_0_, *y*_0_, *x*_0_) into current study data *D* = (*n, y, x*). A parameter *a*_0_ *∈* [0, 1] controls the influence of the historical data on the prior distribution. The use of power prior in psychophysical data was described in detail in our previous article (Mezzetti et al., 2023).

The Bayesian model in Eq. (15)–(19) can be extended to include guess and lapse rates, in analogy with Eq. (8) (BH-GNM). Accordingly, Eq. (15) at the first level is modified as:

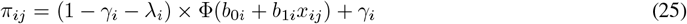

Similarly, it is possible to extend the model parameterized as in Eqs. (20)–(24). As for the GNM, we approximated *pse*_*i*_ = *µ*_*i*_ and *jnd*_*i*_(*p*) = *σ*_*i*_[Φ^*−*1^(*p*) *−* Φ^*−*1^(*−* 0.5)]. To include lapse and guess rates as random effects, the following prior distributions are specified:

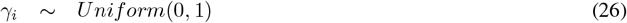

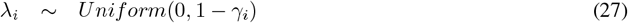

The condition in Eq. (27) guarantees that *γ*_*i*_ + *λ*_*i*_ *<* 1 for every *i*.

### Model Diagnostics and Statistical Inference

Assessing a model fit is essential for evaluating the validity of the assumptions and driving model selection. The first step should always involve visual inspection of the fitted model. Here, we relied on visual evaluation for the model fits to individual participants. We elaborated on this rationale in the Discussion section.

Additional diagnostics can support model evaluation and selection, including analysis of the residuals (Santos Nobre & da Motta Singer, 2007) by means of plots. Furthermore, model comparison was based on the sum of squared errors (SSE), which quantifies the overall deviation between model predictions and observed data. SSE offers a direct and framework-independent measure of goodness of fit, allowing fair comparison between frequentist and Bayesian models. For nested frequentist mixed models, information criteria such as AIC or BIC can help determine which effects to include (Müller et al., 2013). Conversely, likelihood-based criteria such as AIC or BIC cannot be employed to compare frequentist and Bayesian models, as the likelihoods of frequentist and Bayesian models are not directly comparable due to differences in parameter treatment and prior assumptions.

All Bayesian models were estimated using Markov Chain Monte Carlo (MCMC) sampling (Gilks et al., 1995). As is standard practice, convergence diagnostics need to be performed to verify chain stability and effective sampling, ensuring reliable posterior inference. For chain diagnostics, the most straightforward and widely used method is the 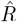 statistic (Gelman & Rubin, 1992; Vehtari et al., 2021). The 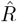 statistic compares the between-chain and within-chain variances to assess convergence; values close to 1 indicate that the chains have likely converged, while larger values suggest non-convergence. In this study, 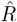 was used to confirm model convergence, followed by graphical inspection of the posterior chains, often supported by trace plots for visual confirmation. Different R packages (i.e. *bayesplot* (Gabry *et al*., 2019), *coda* (Plummer et al., 2006), *boa* (Smith, 2007)) can be used to monitor the convergence of MCMC chains. In this study, we used *ShinyStan* (Muth et al., 2018), an interactive R package for the visualization and diagnostics of Bayesian model samples. *ShinyStan* provides trace plots, posterior density estimates, autocorrelation plots, and pairwise parameter comparisons, enabling evaluation of chain convergence, 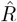 effective sample size, and potential model issues.

Although direct comparison between frequentist and Bayesian models is not straightforward, model diagnostics remain valuable tools for assessing fit and guiding model selection. In this paper, along with classical model diagnostics based on residuals, we illustrate the use of simulated-residual tests, implemented in R with the *DHARMa* package (Hartig, 2024), to evaluate hierarchical models in both frequentist and Bayesian frameworks.

In our examples, the focus of statistical inference differs by dataset: in the first, we assess the model fit by comparing estimated parameters to the simulated (or expected) values, whereas in the second, we evaluate experimental effects by comparing parameter estimates across conditions. For the single-subject models, statistical inference is carried out through a second-level analysis (in our case, one-sample or paired t-tests) applied to fitted or derived parameters. Although it is frequently used in psychophysical literature, this procedure is problematic because it ignores the standard errors associated with each individual estimate. In GLMMs, inference is incorporated into the model via the population-level (fixed) effects, with uncertainty on the estimated parameters quantified through bootstrap procedures (Moscatelli et al., 2012). For Bayesian models, inference consists in inspecting the posterior distribution of the model parameters (McElreath, 2018; Mezzetti et al., 2023).

### Software

In this tutorial, we demonstrate the use of various R tools and packages for data preprocessing, modeling, and visualization. To follow along with the examples, we recommend that the reader have an R environment running with the required packages installed. The R code in this tutorial was tested on Mac, Ubuntu, and Windows operating systems, using multiple R releases and the RStudio IDE. A detailed report of the R environment and package versions is available in the Supplementary Information. The script in R and Stan are explained in this tutorial and available for download in the following Github repository: github.com/moskante/hierarchical-models-psychophysics.

### Preprocessing, data wrangling, and visualization

For data preprocessing and visualization, we rely on packages from the *tidyverse* suite (Wickham & Grolemund, 2016), which includes several packages that support efficient and readable data workflows. In particular, *dplyr* is used for data manipulation and *purrr* allows for functional programming. The *ggplot2* package is used for creating visualizations, and *magrittr* enables piping operations through the `%>%` (pipe) operator.

### GLM

To fit the model in Eq. (2) to each individual participant’s data, we use the base R function `glm()` from the *stats* package (loaded by default in R). Additional utilities are provided by our own package, *MixedPsy*, which simplifies common GLM-based psychophysical analyses.

### GNM

To fit the model in Eq. (8) using the GNM framework, we use the function `gnlr()` from the *gnlm* package (Yssaad-Fesselier & Knoblauch, 2006; Swihart & Lindsey, 2025). Custom functions for model fitting and bootstrapping are provided and made available for download.

### GLMM

We use the *lme4* package to fit the GLMMs in Eq. (11) (Bates et al., 2015). The *lmerTest* package (Kuznetsova et al., 2017) augments *lme4* by providing p-values and F-tests for fixed effects through Satterthwaite/Kenward–Roger approximations (Halekoh & Højsgaard, 2014). *DHARMa* offers tools for model diagnostic (Hartig, 2024). The *MixedPsy* package provides functions for psychophysical data analysis.

### BHF

The models presented in this tutorial are implemented in Stan, a probabilistic programming language for Bayesian inference (Carpenter et al., 2017). The corresponding `.stan` model files are available for download. We use the *rstan* package to interface with Stan from within R, allowing model compilation and sampling (Stan Development Team, 2025). For diagnostic checks and model visualization, we employ functions from *DHARMa* and the *shinystan* package (Gabry & Veen, 2022).

## Results

### Example 1: A simulated dataset

#### GLM

To fit a GLM for a single participant in `simul_data`, the R code is the following:

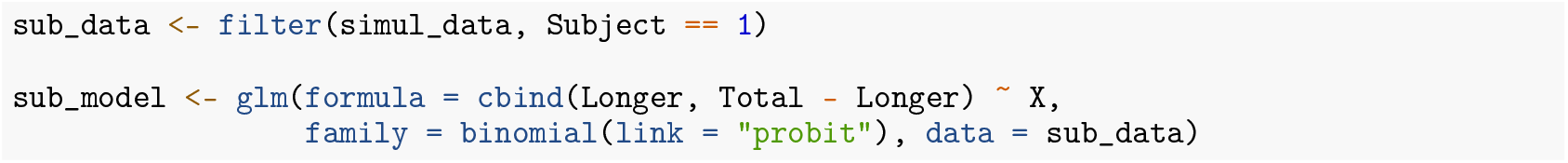

In accordance with Eq. (2), the `formula` argument defines the relationship between the response and predictor variables, with the binomial response on the left and the continuous predictor (`X`) on the right of the tilde symbol (~). The response variable is structured as a two-column matrix, where each column represents the number of successes and failures, respectively, combined using `cbind()`. The `family` argument specifies the error distribution of the response variable (binomial) and the link function (probit). Alternative link functions for binomial data in `glm()` include `logit`, `cauchit`, and `cloglog`. These correspond to cumulative distribution functions (CDFs) for logistic, Cauchy, and to the complementary log-log link function, respectively.

An overview of the fitted model, including the intercept and slope parameters (i.e., *β*_0*i*_ and *β*_1*i*_ from Eq. (2)), can be obtained by calling `summary(sub_model)`. The parameters of the psychometric function can be extracted by applying Eq. (3) and (4) to the model coefficients, which are accessed via `coef(sub_model)`. Since PSE and JND are non-linear combinations of the model parameters, their variance cannot be directly estimated from the standard errors of the model coefficients. In our previous work (Moscatelli et al., 2012) we demonstrated how to estimate PSE and JND along with their variance using either the delta method (Casella & Berger, 2002) or the bootstrap method (James et al., 2013) in GLM and GLMM framework. The delta method is based on the assumption that the estimator asymptotically follows a normal distribution, and approximates the mean and variance of a function of random variables using its truncated Taylor series expansion (Casella & Berger, 2002). The package *MixedPsy* includes the function `PsychDelta(sub_model)` which computes PSE and JND along with their variance using the delta method for an object of class `glm` with a probit link function.

To estimate psychometric parameters for all participants in an experiment, the steps described above must be applied to each participant individually, and the resulting parameters compiled into a data frame for second-level analysis. This can be done in R using various approaches, such as the base function `apply()` or the `map()` function from the *purrr* package, both of which allow for efficient iteration over multiple participants. The *MixedPsy* package provides convenient wrapper functions built on the *purrr* framework, enabling users to fit models across subsets of data and automatically generate a well-structured data frame of estimated parameters:

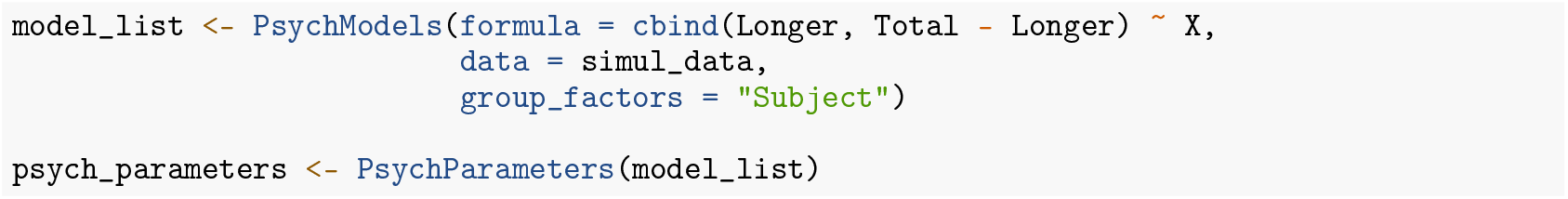

In addition to storing the psychometric parameters in a data frame for further analysis, it is often necessary to visualize the model, to evaluate or report results. For a GLM such as `sub_mod`, this can be done by plotting the model’s predicted values using:

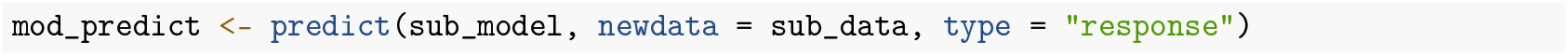

Since the link function in GLMs is generally not the identity, the relationship between predictors and the response mean is non-linear, requiring interpolation of the independent variable to produce a smooth prediction curve. The *MixedPsy* package provides the `PsychInterpolate()` function, which automates the process by interpolating and generating predictions for a list of fitted psychometric models:

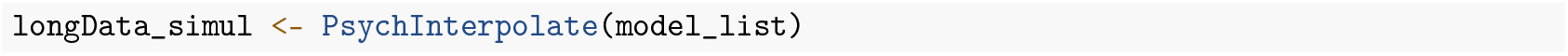

The model predictions, together with the data, can be visualized using *ggplot2*:

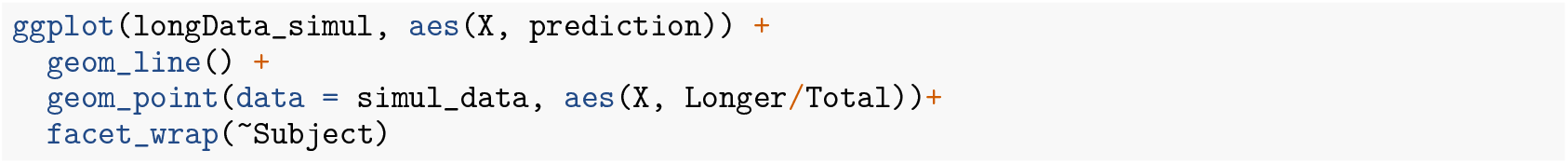

The resulting plots are shown in Fig. 1 (red lines).

**Figure 1.**
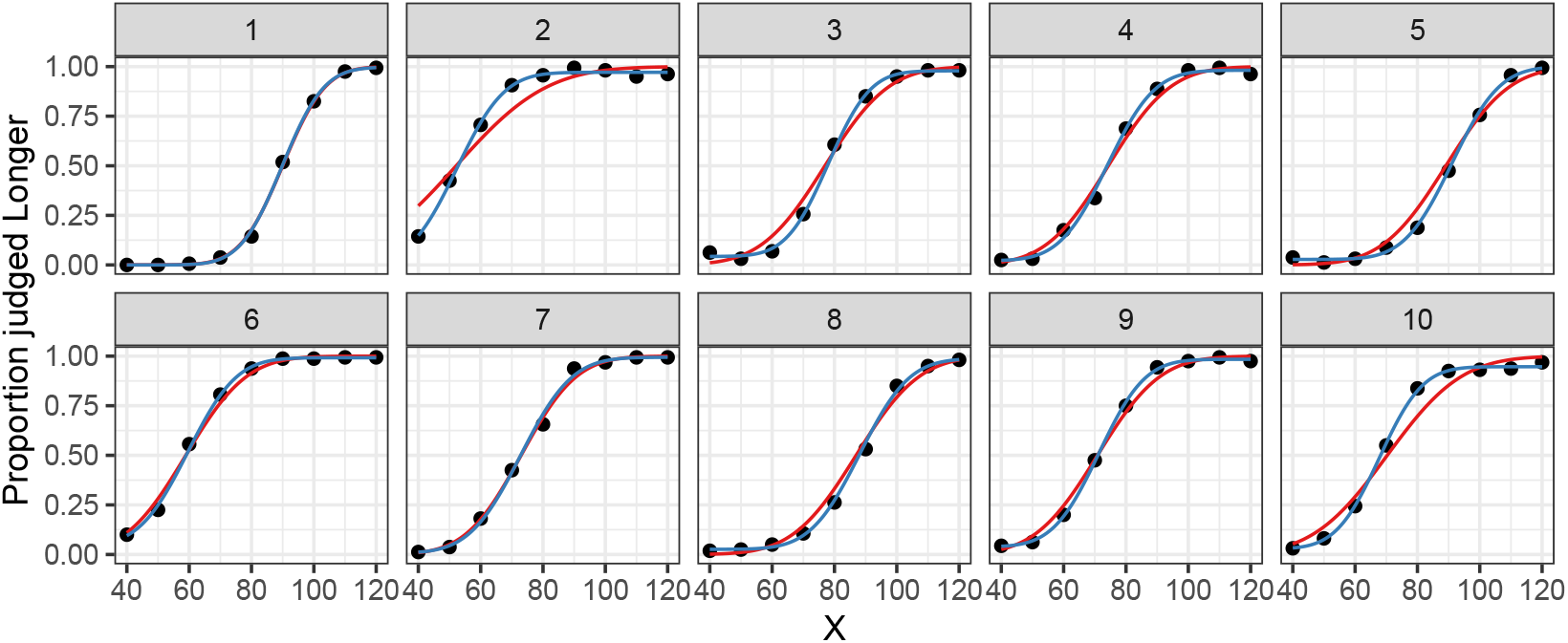
Single-subject model predictions for simulated data: GLM (red lines), GNM (blue lines), with simulated proportions shown as black points.

#### GNM

To fit the model in Eq. (8) using `gnlr()`, the required inputs include a response vector (or matrix for successes and failure, as in the GLM example above), a predictor variable vector, a link function, and initial parameter values. The *gnlm* package does not accept data frames as input, therefore the data must be provided explicitly as separate arrays:

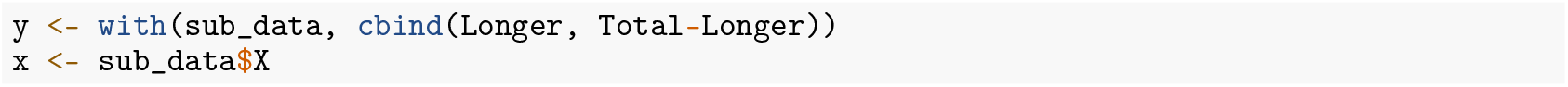

The link function can be defined to match the relationship in Eq. (8). To ensure valid probability estimates and avoid fitting errors, it is essential that the guess and lapse rates remain within specific bounds. Specifically, for each participant *i, γ*_*i*_ must lie between 0 and 1, while *λ*_*i*_ must lie between 0 and 1 *− γ*_*i*_.

Since the *gnlm* package does not allow direct specification of parameter constraints, a common workaround is to apply transformation functions that implicitly enforce these bounds. Following Yssaad-Fesselier & Knoblauch (2006), the arctangent function can be used to constrain the parameters as follows:

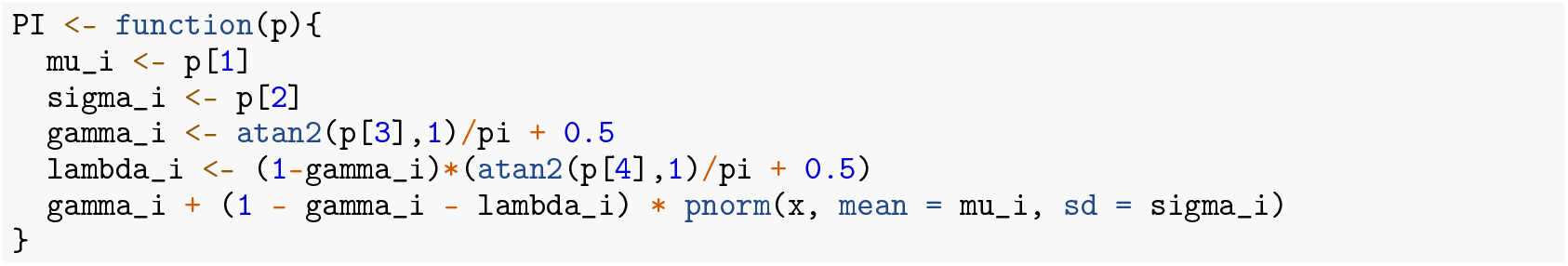

To aid model convergence, it is important to provide reasonable starting values for the parameters. Suitable initial estimates for *µ*_*i*_ and *σ*_*i*_ can be obtained by first fitting a GLM, as illustrated in the previous section. For *γ*_*i*_ and *λ*_*i*_, a practical first approximation is to use the observed proportions of responses at the lowest and highest stimulus levels, respectively. Custom function `pstart()` computes starting values while respecting the parameter constraints and transformations applied to *γ*_*i*_ and *λ*_*i*_ in the definition of the link function `mu()`.

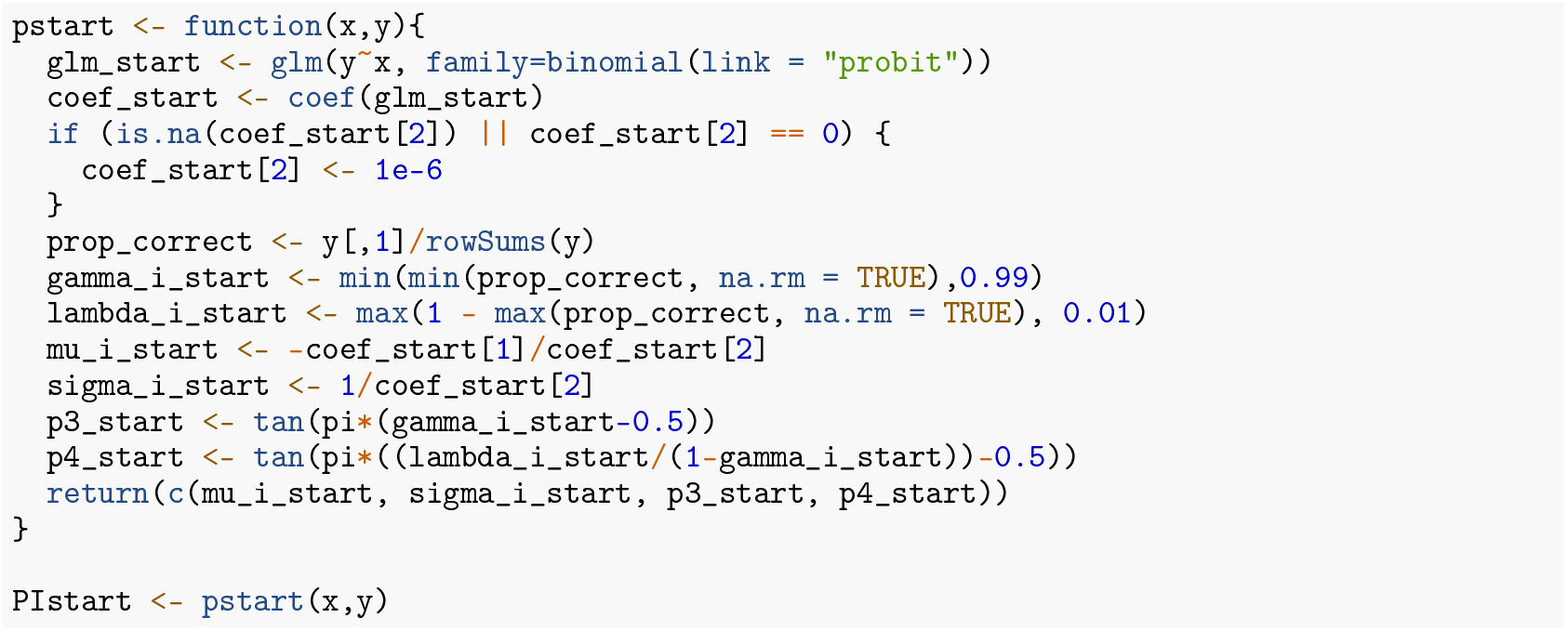

The model can then be fitted with `gnlr` as follows:

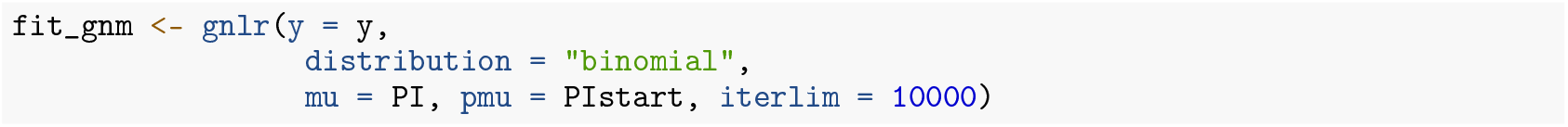

The parameters of the psychometric function can be recovered from the fitted model using the custom function `PsychParametersGNM()`, which accounts for the parameter transformations applied in model specification:

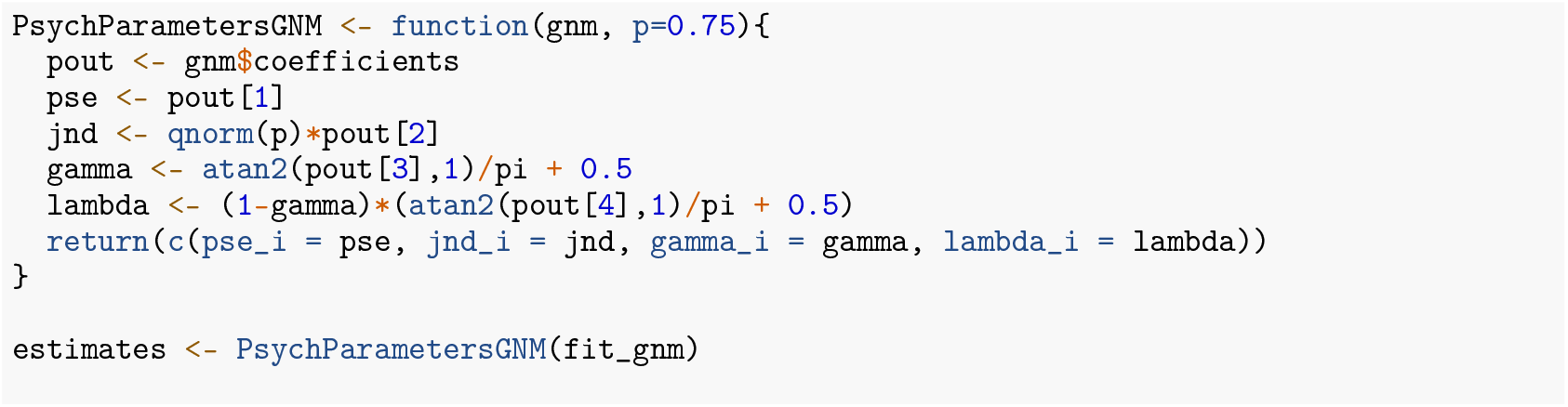

The variance of the estimated parameters can be assessed with a bootstrap procedure via the *boot* package (Angelo Canty & B. D. Ripley, 2024). As with GLMs, the model-fitting pipeline is applied to each participant individually to obtain their parameter estimates. Example functions demonstrating the process are available in the repository associated with this publication. The resulting fits are shown in Fig. 1 (blue lines).

The functions described above can be customized to suit specific experimental or modeling needs. For instance, the lapse and guess rates can be omitted entirely or fixed to predetermined values—commonly, the guess rate is fixed at 1/n in n-AFC tasks. Additionally, the normal CDF used in the model can be replaced with alternative functions such as the logistic or Weibull CDF, depending on the theoretical assumptions or empirical fit of the data.

#### GLMM

The following code fits the GLMM described in Eq. (10) to `simul_data`:

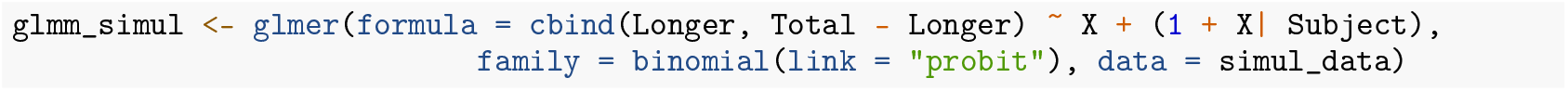

As in `glm()`, the relationship between the response variable and the linear predictor is defined using the formula argument, and the error distribution and link function are specified via the `family` argument. In the formula, terms outside the parentheses represent fixed effects, while terms inside the parentheses define the structure of the random effects and the clustering variable (indicated after the `|` symbol). In the example, a random intercept and a random slope are included for each subject, along with their covariance. By default, the covariance between random effects is estimated unless explicitly excluded using the || syntax–e.g., `(1 + X || Subject)`.

The package *MixedPsy* provides functions for estimating psychometric parameters. For easier maintenance and compatibility, most *MixedPsy* functions that rely on the *lme4* package do not operate directly on model objects of class `glmerMod`. Instead, they require an intermediate object of class `xplode`, which can be created as follows:

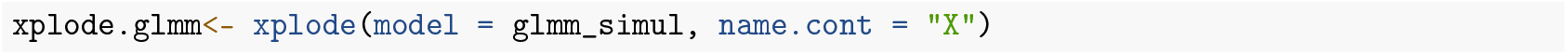

Once the model is converted to an `xplode` object, the `MixDelta()` function can be used to compute estimate and 95% confidence intervals of PSE and JND using the delta method (Casella & Berger, 2002):

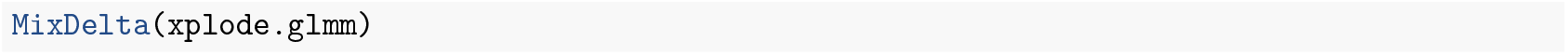

Because the delta method assumes asymptotic normality of the parameters estimates, it may be not appropriate in the context of GLMMs. Therefore, results obtained with `MixDelta()` should be used as preliminary approximations, while a bootstrap approach is recommended for more robust and accurate estimation. This can be implemented using the `bootMer` function from the *lme4* package, in combination with user-defined functions for extracting PSE and JND. The `pseMer()` function from *MixedPsy* provides a wrapper for performing bootstrap resampling directly on a `glmerMod` object.

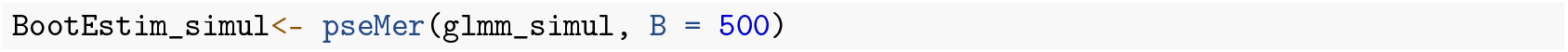

For GLMMs with a single continuous predictor, `pseMer()` provides by default the bootstrap estimates of the sample’s PSE and JND. Increasing the number of bootstrap iterations via the `B` argument typically improves the precision of the estimates but also increases computational time.

The *MixedPsy* package provides the `MixInterpolate()` function to stremaline the process of interpolating a fitted GLMM over a grid of predictor values for visualization (Fig. 2, orange lines).

**Figure 2.**
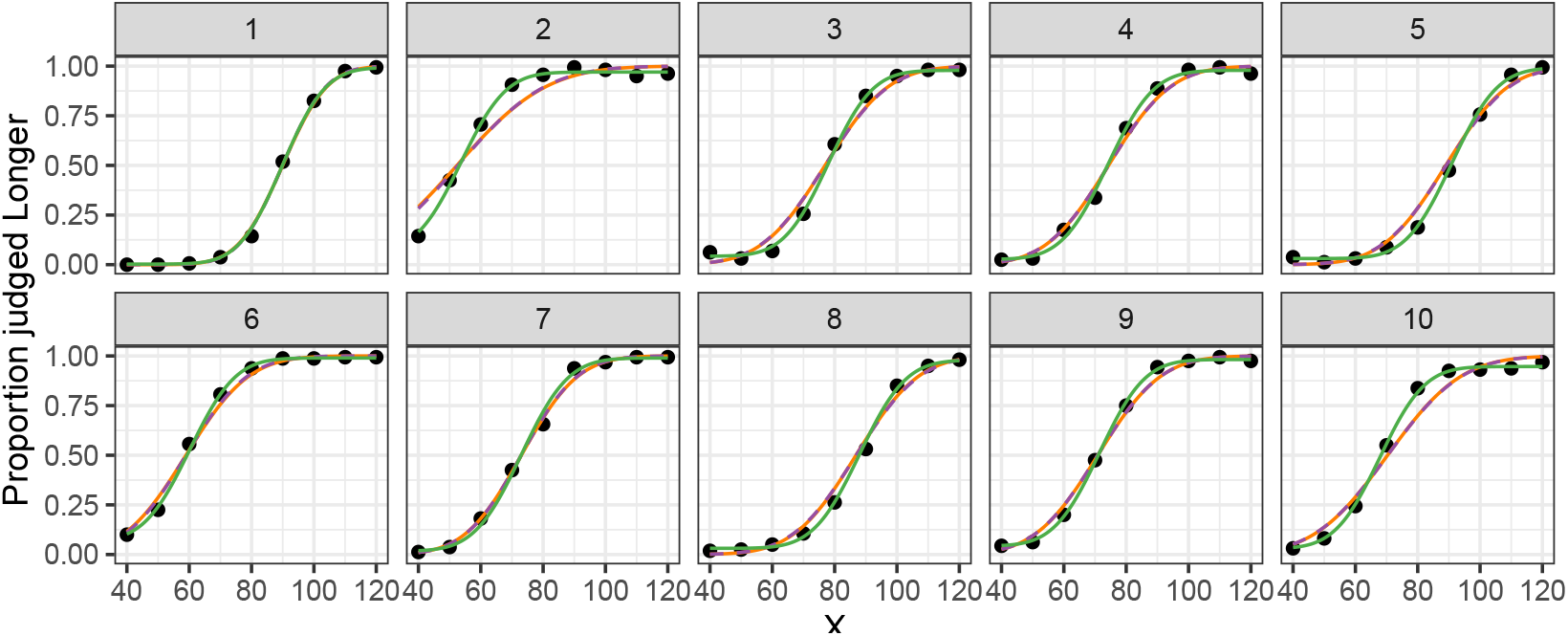
Hierarchical model predictions for simulated data: GLMM (orange lines), BHF without lapse and guess rates (purple lines), and BHF with subject-specific lapse and guess rates (green lines). Simulated proportions are shown as black points.

##### BHFs

Here, we show how to fit the model in Eq. (20)–(24) with Stan.

A Stan program consists of several blocks. The `data` block declares the observed variables required by the model:

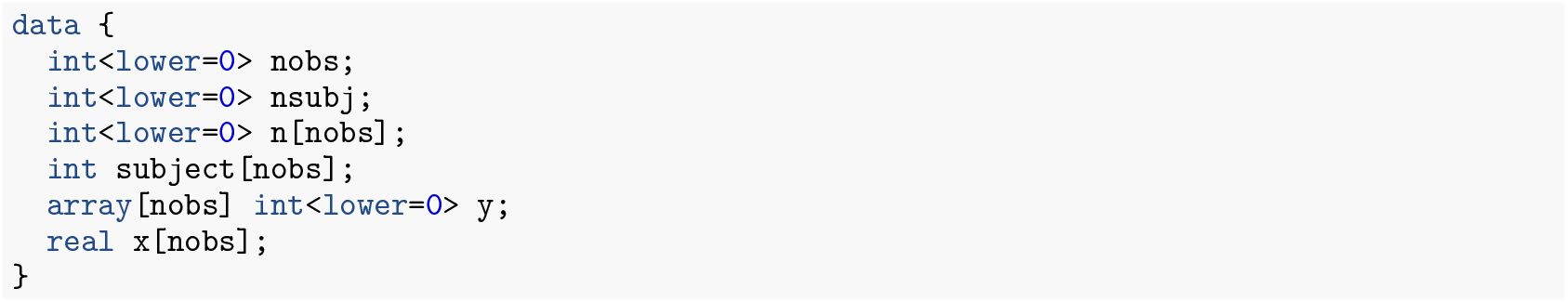

The `parameters` block defines the model parameters and hyperparameters that Stan will estimate, including their boundaries and constraints. For the BH-GLM described above, using the parameterization in Eq. (20), the parameters are specified as follows:

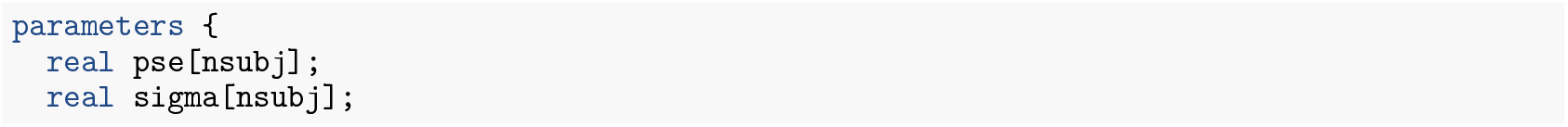

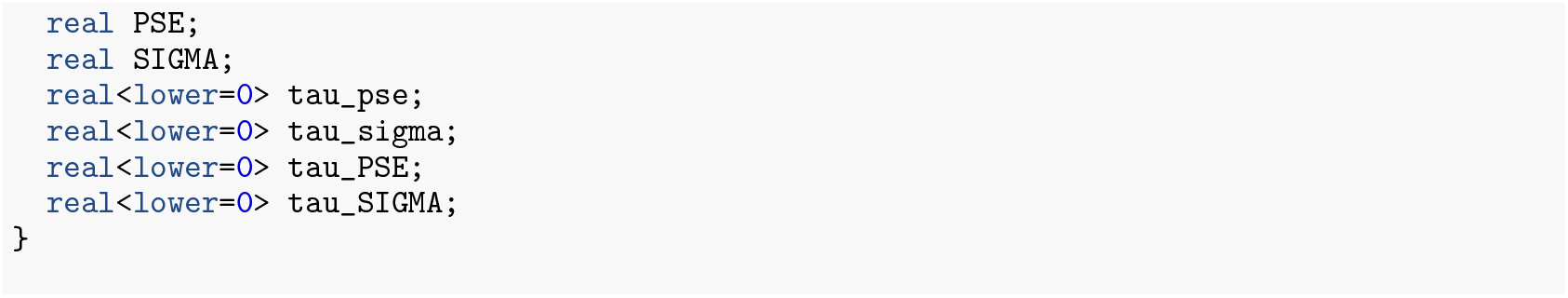

The `transformed parameters` block is used to define of intermediate quantities and optional variables. Variables declared in this block are included in the model output. In this example, we use the block to compute Φ^*−*1^(*π*_*ij*_), as defined in Eq. (20):

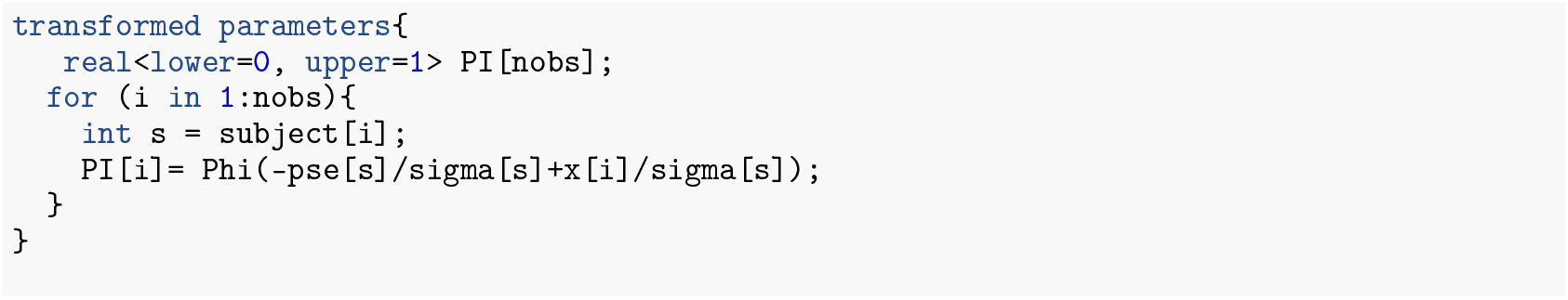

Constraints on the transformed parameters, such as the bounds on `pi1`, are validated after the statements are executed.

Since the model estimates both population-level and subject-level of sigma, labeled as Σ and *σ*_*i*_, respectively, the block can be used to define population-level and individual JNDs:

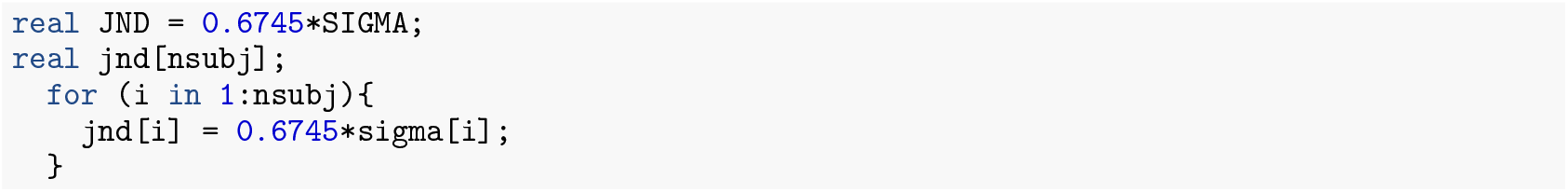

The `model` block specifies the prior distributions, hierarchical structure, and likelihood. Note that the variables declared here are local to the block and excluded from the model output, unless explicitly declared elsewhere. The following block implements the model in Eq. (1), with parameters defined as in Eqs. (20)–(24), and with hyperprior distributions on the variance components drawn from Cauchy distributions:

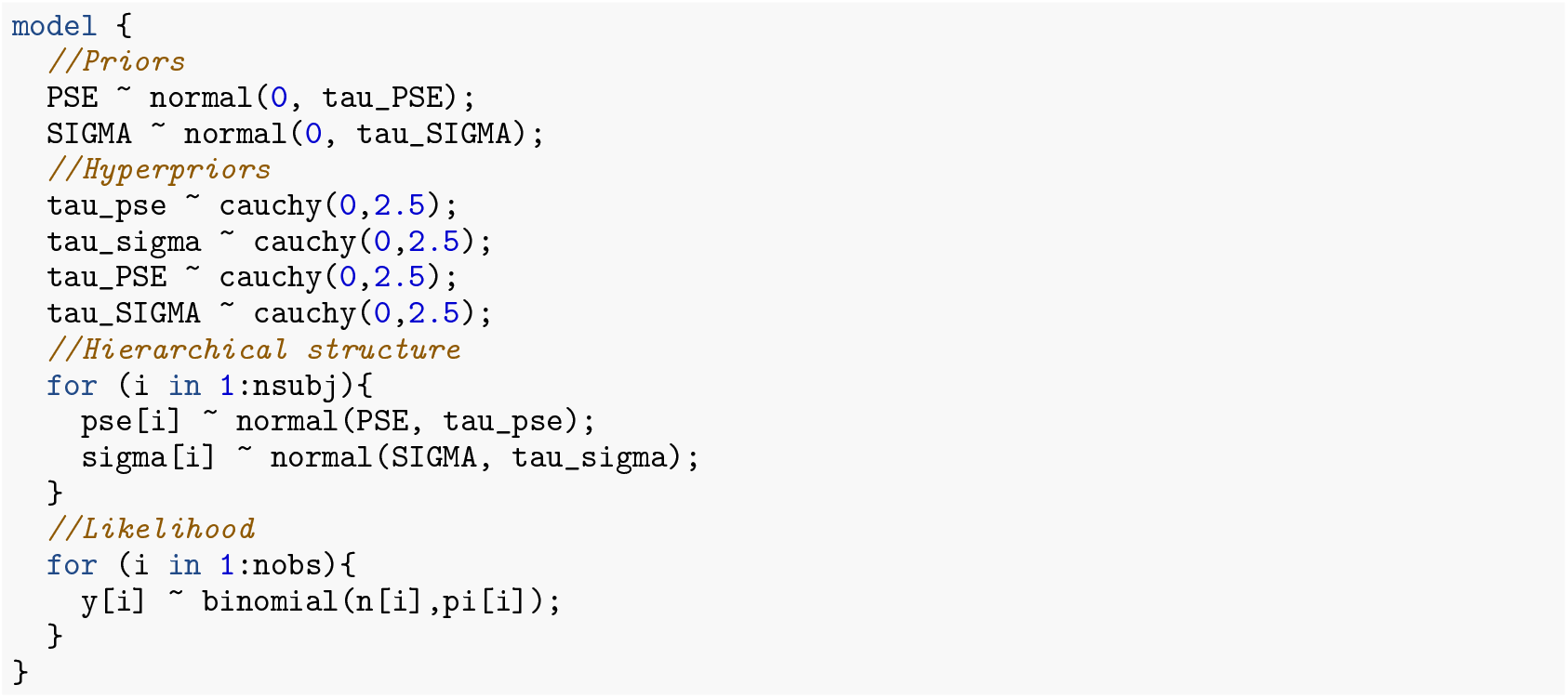

Noninformative priors have been used in this code. Refer to Mezzetti et al. (2023) for examples of models with informative priors.

For model diagnostics, the package *DHARMa* requires that the Stan model includes a `generated quantities` block that produces the necessary posterior predictive simulations and predicted response values. These quantities are essential for DHARMa to compute simulated residuals and perform model diagnostics.

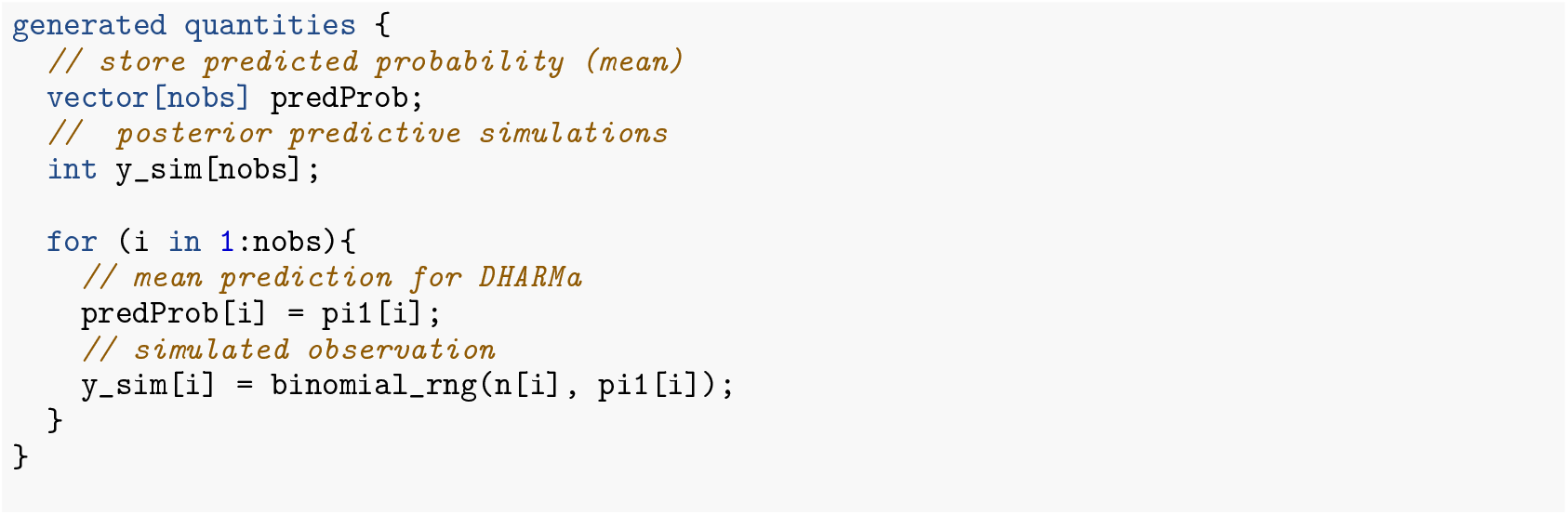

The Stan program is compiled and executed from R using the `stan()` function in the *rstan* package. The function requires the path to the Stan file and a named R list whose elements match the variables of the `data` block in the Stan program. Other specifications—such as the number of Markov chains, total iterations, and the number of warmup iterations—can also be specified within the `stan()` call.

By default, Stan initializes all parameters with random values between −2 and 2 on the unconstrained scale. However, custom initial values can be provided via the `init` argument. In our examples, we initialized the parameters `pse` and `sigma` using the function below and fitted the Stan model as follows:

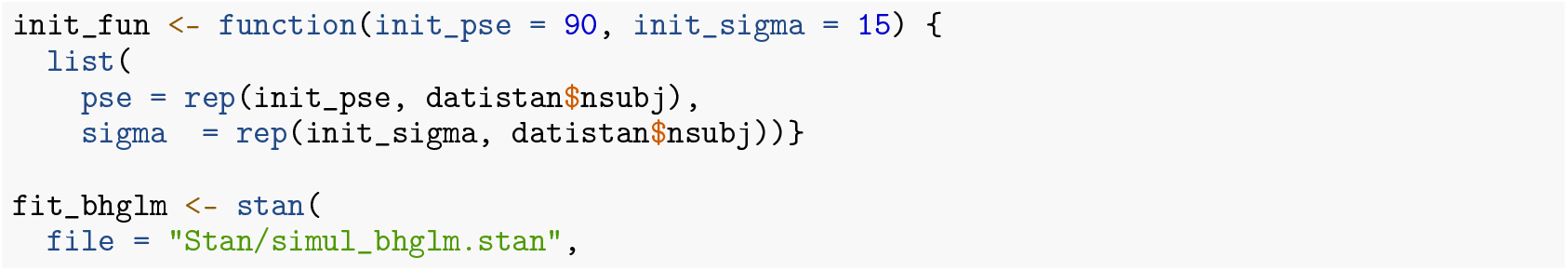

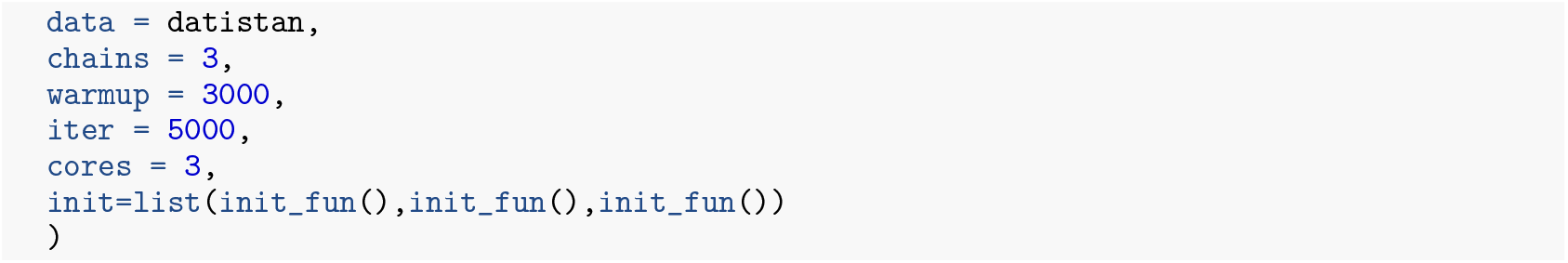

The posterior estimates of the parameters, computed as the means of the posterior samples, can be used to generate and visualize model predictions, as illustrated in Fig. 2 (purple lines).

To extend the model to include lapse and guess rates (BH-GNM), the response probability must be adjusted accordingly. This requires adding the guess and lapse parameters, *γ*_*i*_ and *λ*_*i*_, to the `parameters` block:

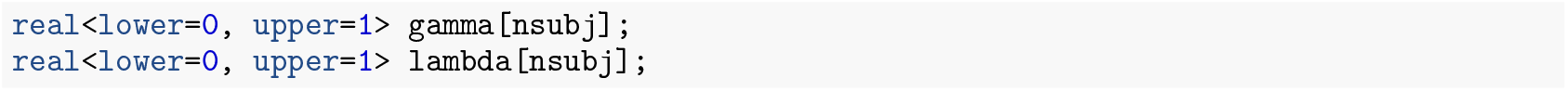

The structure of the response probability must be modified according to Eq. (25):

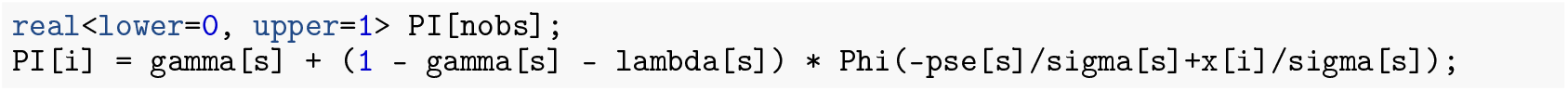

Finally, the priors for *γ*_*i*_ and *λ* _*i*_ (Eqs. (26)–(27)) are specified in the model block:

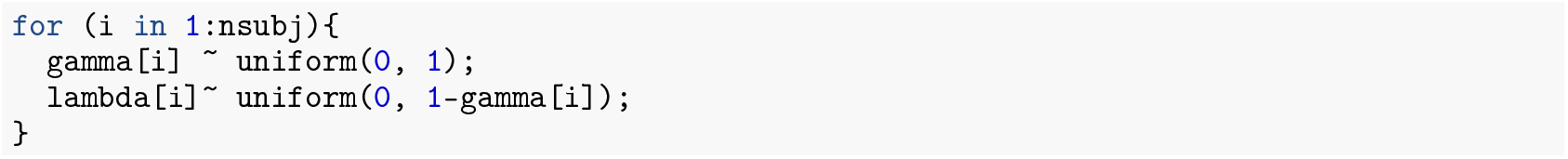

The `stan` function is called to compile and run the Stan program as previously described. In the parameter initialization function, we added the lapse and guess rates, ensuring they are initialized to positive values:

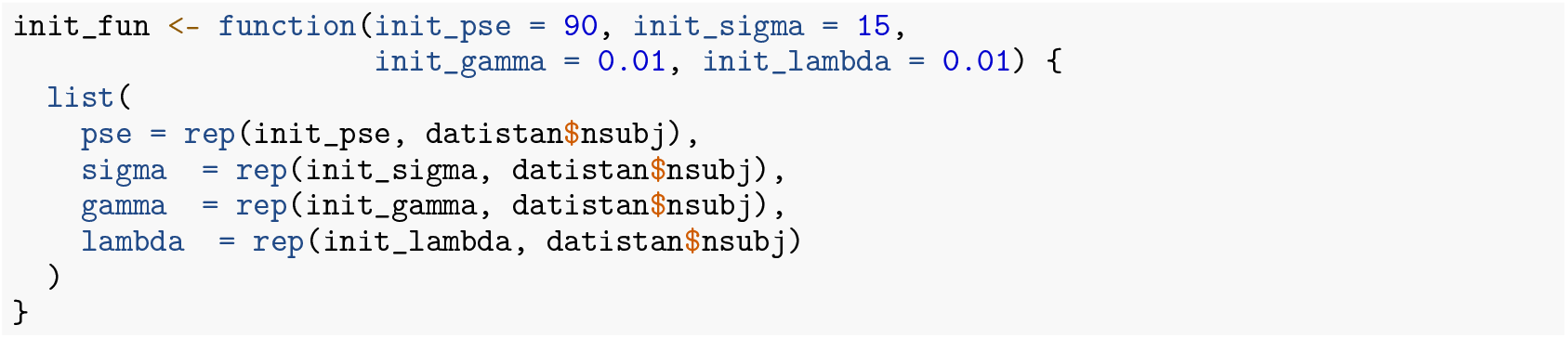

Model predictions for the BH-GNM are shown as green lines in Fig. 2.

### Model Diagnostics and Statistical Inference

From visual inspection of single-subject models (Fig. 1), it appears that for several participants the GLM predictions do not align well with the observed data. This discrepancy is expected, as the simulated dataset includes small but non-zero guess and lapse rates that affect behavior at low and high stimulus levels. Accordingly, the GNMs more accurately capture behavior at the asymptotes of the psychometric functions. Similar patterns are observed for the hierarchical models (Fig. 2), where the GLMM and BH-GLM yield nearly identical predictions and generally provide a poorer fit compared to the BH-GNM. The SSE values for the three hierarchical models are reported in Table 4. As expected, the BH-GNM shows the lowest SSE among the three models, consistent with its larger number of parameters. The reduction in SSE is substantial—approximately an order-of-magnitude improvement—reflecting the markedly better fit provided by the model.

**Table 4.**
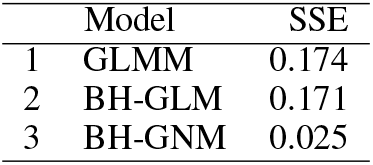
SSE for residuals of hierarchical models in Example 1.

|  | Model | SSE |
| --- | --- | --- |
| 1 | GLMM | 0.174 |
| 2 | BH-GLM | 0.171 |
| 3 | BH-GNM | 0.025 |

Diagnostic checks using DHARMa are presented in Supplementary Fig. 1. Tests on the simulated residuals show significant quantile deviations for both the GLMM and BH-GLM (top and middle panel, respectively), as well as significant deviations from uniformity in the BH-GLM residuals (bottom panel). For the BH-GNM, the residuals show small but statistically significant deviations from uniformity, while no issues are present in the quantile deviations. Overall, DHARMa diagnostic results indicate that models incorporating guess and lapse rates provide a more appropriate fit, both at the single-subject and hierarchical level.

Using the *Shinystan* package confirms the stability of the sampling chains and, consequently, of the posterior estimates. The values for the 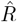 statistics are below 1.02 for all the parameters in both the BH-GLM and BH-GNM models, although the latter requires a larger number of iterations (15000 iterations after 10000 burn in). Running BH-GNM model a negligible number of sampling warnings after warm-up were detected.

Next, we evaluated the estimated values of PSE and JND across the different models (Figs. 3 and 4). PSE estimates were close to the true value (black bars in the figure) across all models and for all the simulated observers (Fig. 3, top panel). The bottom panel in Figure 3 shows the estimates of the five model at population level. The true mean of PSE = 74.6 was labeled as a black solid line in the figure. We assessed whether the estimated means differed from the true mean used in the simulations. In the GLM and GNM frameworks this required a two level approach, and one-sample t-tests for the GLM and GNM showed no significant differences with the true value of PSE of 74.6 (all t < 0.1, all p > 0.9). A boostrap proceduere was used for the GLMM and the bootstrap-based 95% confidence intervals included the true value of PSE (CI = [67.5, 81.9]). For both Bayesian models, 95% credible intervals contained the true value of PSE (BH-GLM CrI = [65.6, 82.4], BH-GNM CrI = [65.3, 82.2]).

**Figure 3.**
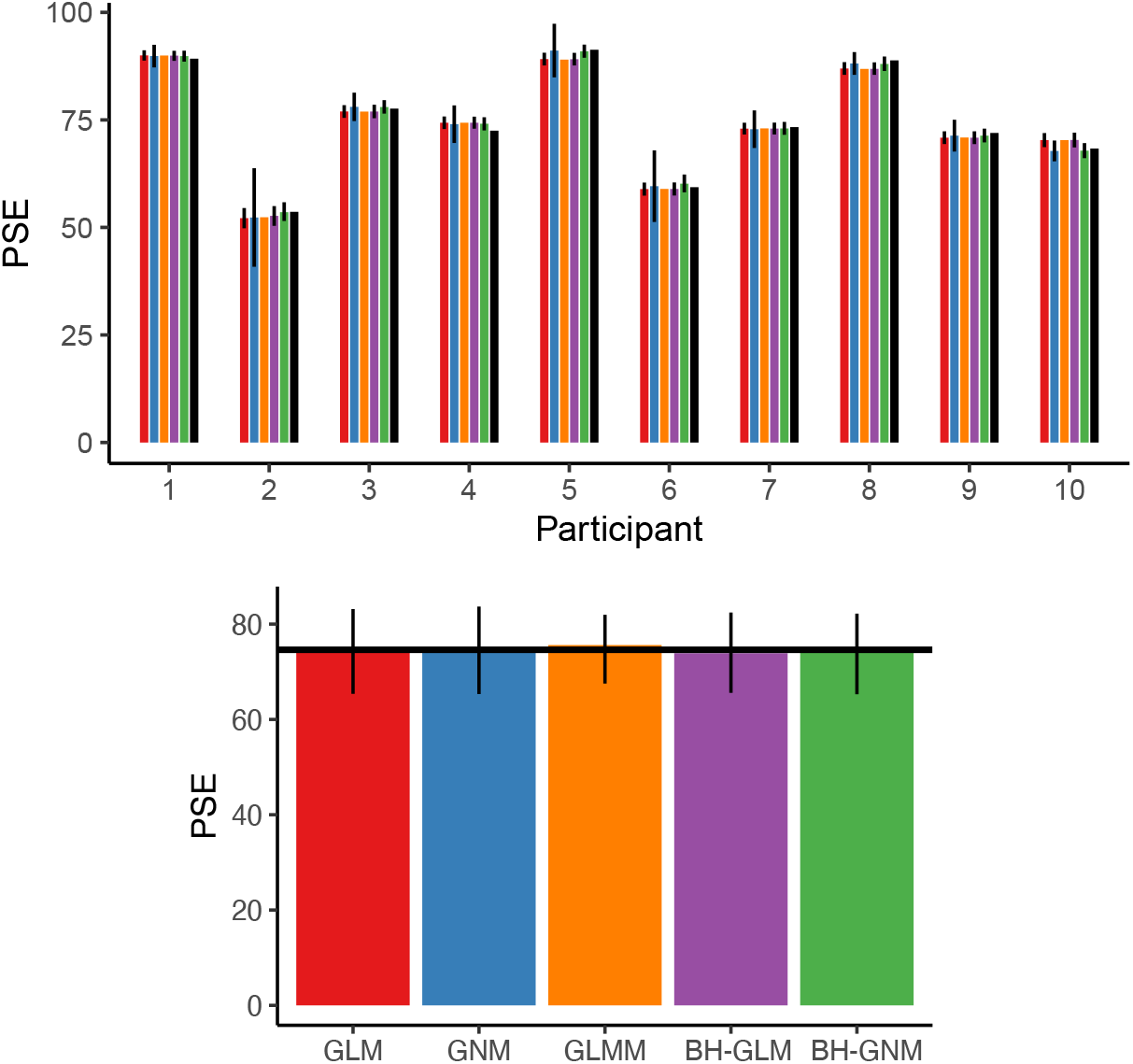
PSE estimates for each simulated participant in the dataset (top) and from the simulated dataset (bottom) across the 5 models. Colors indicate the model used: red = GLM, blue = GNM, orange = GLMM, purple = BH-GLM, green = BH-GNM. Top: Black bars represent the true values of PSE for each participant from the simulations. Error bars show 95% uncertainty intervals: confidence intervals based on Delta method for GLM, bootstrap-based intervals for GNM, credible intervals for the BHFs. GLMM does not provide a measure of uncertainty for individual participants. Bottom: Error bars represent 95% uncertainty intervals: sample-based confidence interval for GLM and GNM; bootstrap-based confidence interval (500 iterations) for GLMM; credible intervals for the BHFs. Black horizontal lines represent the mean parameters values of the simulated data. The true value of the PSE is indicated with a solid black line (PSE = 74.6) for the population.

**Figure 4.**
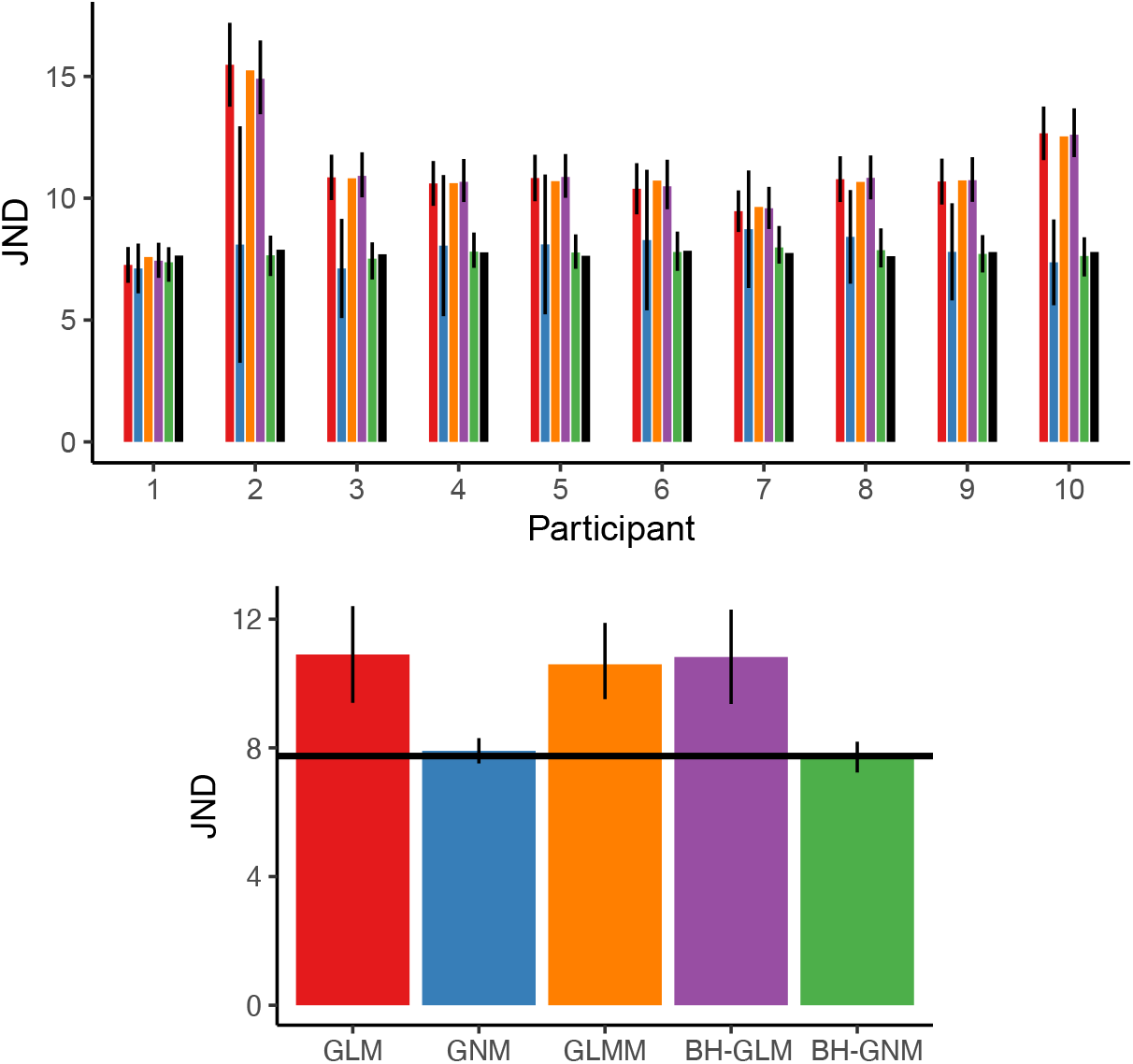
JND estimates for each simulated participant in the dataset (top) and from the simulated population (bottom) across the 5 models. The true values of the JND from the simulations are labeled with black bars for each participant and with a solid black line (JND = 7.7) for the population.

JND estimates were consistent with the true values for the GNM and the BH-GNM, but not for the models that did not allow lapse and guess rates to be estimated (Fig. 4). In those cases, the JND values were systematically overestimated. At the population level (Fig. 4, bottom panel), the t-test for the GLM indicated a significant difference from the true value of the JND of 7.7 (t = 4.75, p = 0.001), whereas the GNM did not (t = 0.94, p = 0.372). For the GLMM, the bootstrap confidence interval did not include the true value of the JND (CI = [9.5, 11.9]). For the BHFs, the credible interval did not include the true value of the JND when asymptotes were fixed (BH-GLM CrI = [9.4, 12.3]), but did include it when random lapse and guess rates were estimated (BH-GNM CrI = [7.2, 8.2]).

### Example 2: The role of vibration in tactile speed perception

#### GLM

The procedure for estimating the parameters of the psychometric functions is the same as that described for the `simul_data` dataset. Due to the inclusion of an additional independent variable, two psychometric functions must be fitted for each participant–one for the control condition and one for the test vibration condition. The code for an individual model is the follow:

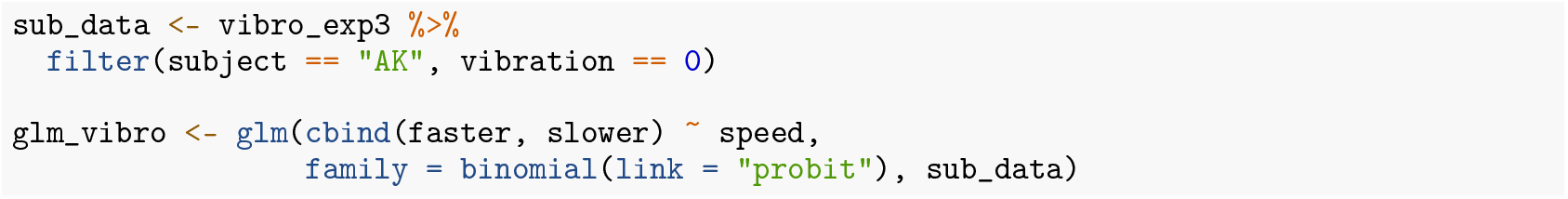

The model can be fitted for all participants and for all vibration conditions using *MixedPsy* functions, by specifying both independent variables as grouping factors when fitting the models:

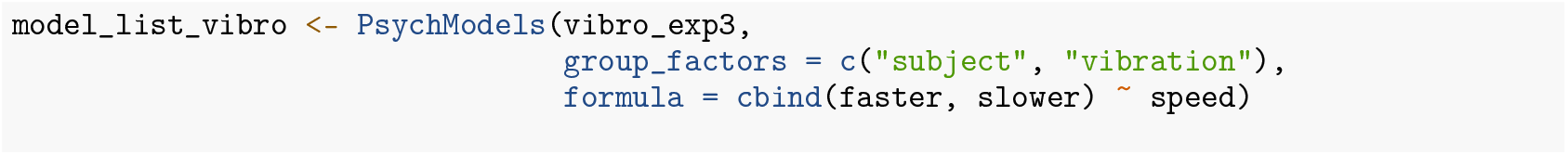

The functions `PsychParameters()` and `PsychInterpolate()` can be applied to the resulting model list to extract psychomentric parameters and interpolate the fitted models for visualization (Fig. 5, red lines). To remain consistent with the original study (Dallmann et al., 2015), we report results in terms of the slope of the fitted models, rather than the JND of the psychometric function. Accordingly, we can obtain the slope estimate and its standard error directly from the fitted GLMs as follows:

**Figure 5.**
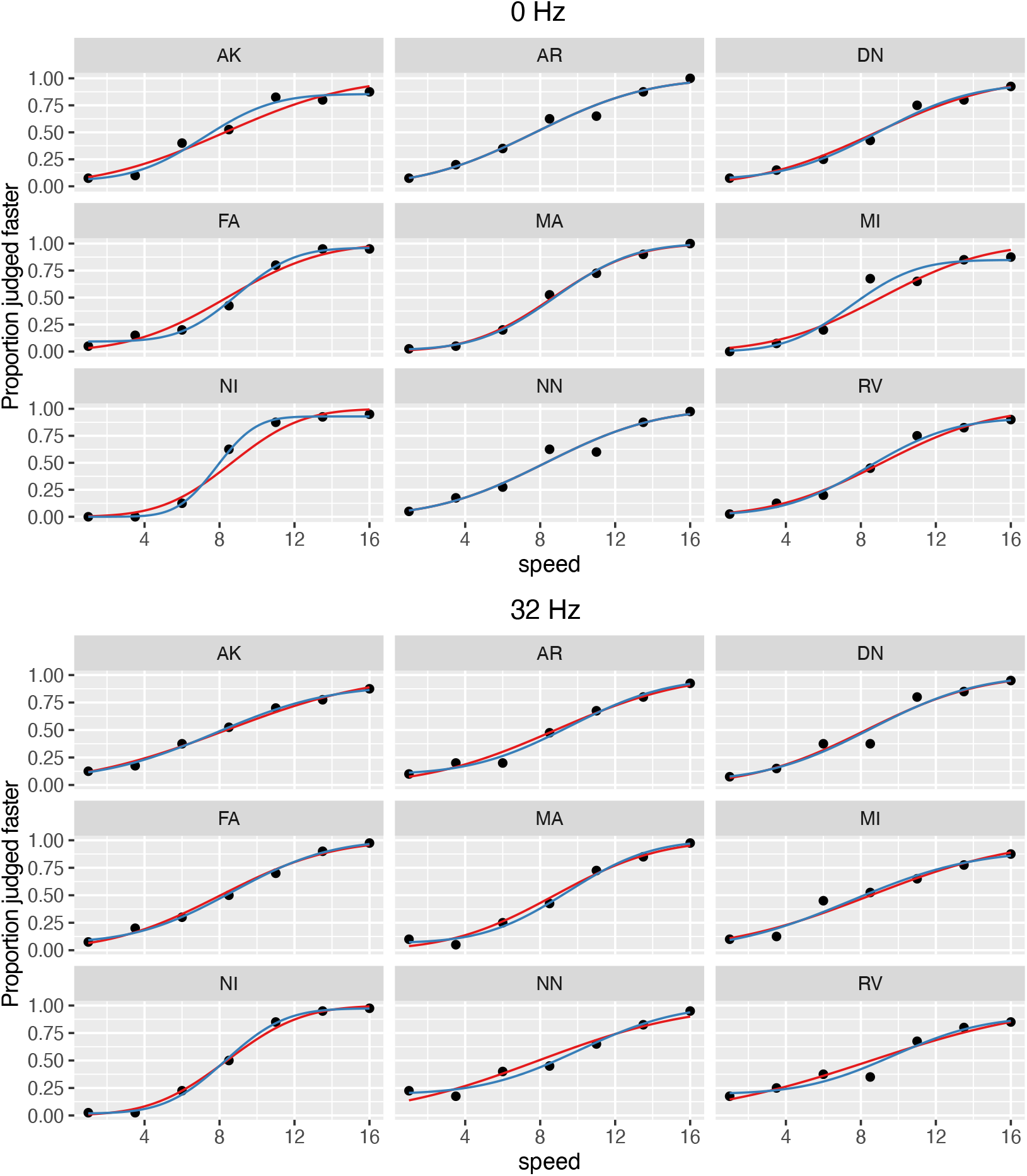
Single-subject model predictions for speed discrimination data: GLM (red lines) and GNM (blue lines), with proportions shown as black points.

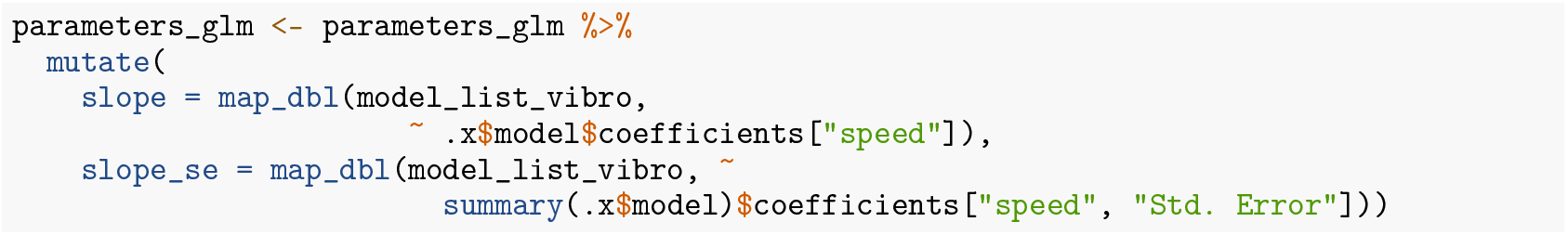

#### GNM

Within the GNM framework, the same model fitting procedure described for the simulated dataset can be applied to the `vibro_exp3` dataset. It is important to include an additional grouping factor to account for the `vibration` condition. Custom functions for fitting GNM models with grouped data are available for download. To obtain slope estimates directly (rather than relying on the reciprocal of the fitted values), the `mu` function and the starting-value function `pstart` can be modified as follows:

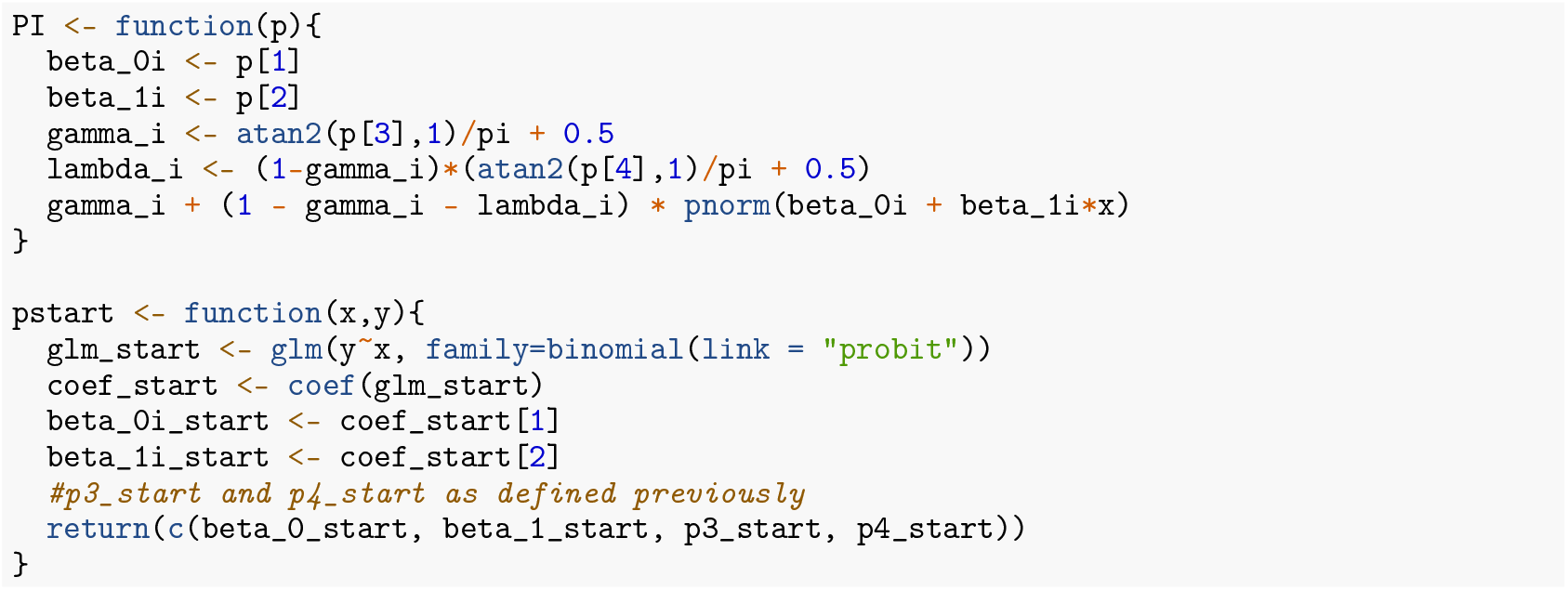

The model fit for individual participants and conditions is shown in Fig. 5 (blue lines).

#### GLMM

For the `vibro_exp3` dataset, we fit a GLMM with the fixed effects specified in Eq. (12) to simultaneously evaluate the population-level effects of speed variation and the presence or absence of masking vibration on the response. The model is implemented using the `glmer()` function as follows:

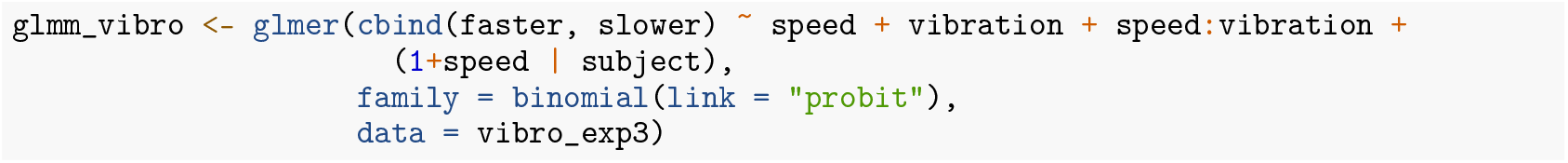

The random-effects structure includes a random intercept, a random slope for speed, and their covariance. This specification was informed by preliminary (not shown) model comparisons. The model coefficients are shown in Table 5, and model fits are shown in Fig. 6 (orange lines).

**Table 5.** Fixed-effect parameters of GLMM for experimental data.

|  | Estimate | Std. Error | z value | Pr(>\|z\|) |
| --- | --- | --- | --- | --- |
| (Intercept) | -2.02 | 0.14 | -14.38 | < 0.001 |
| speed | 0.24 | 0.02 | 14.82 | < 0.001 |
| vibration32 | 0.38 | 0.10 | 3.99 | < 0.001 |
| speed:vibration32 | -0.04 | 0.01 | -4.09 | < 0.001 |

**Figure 6.**
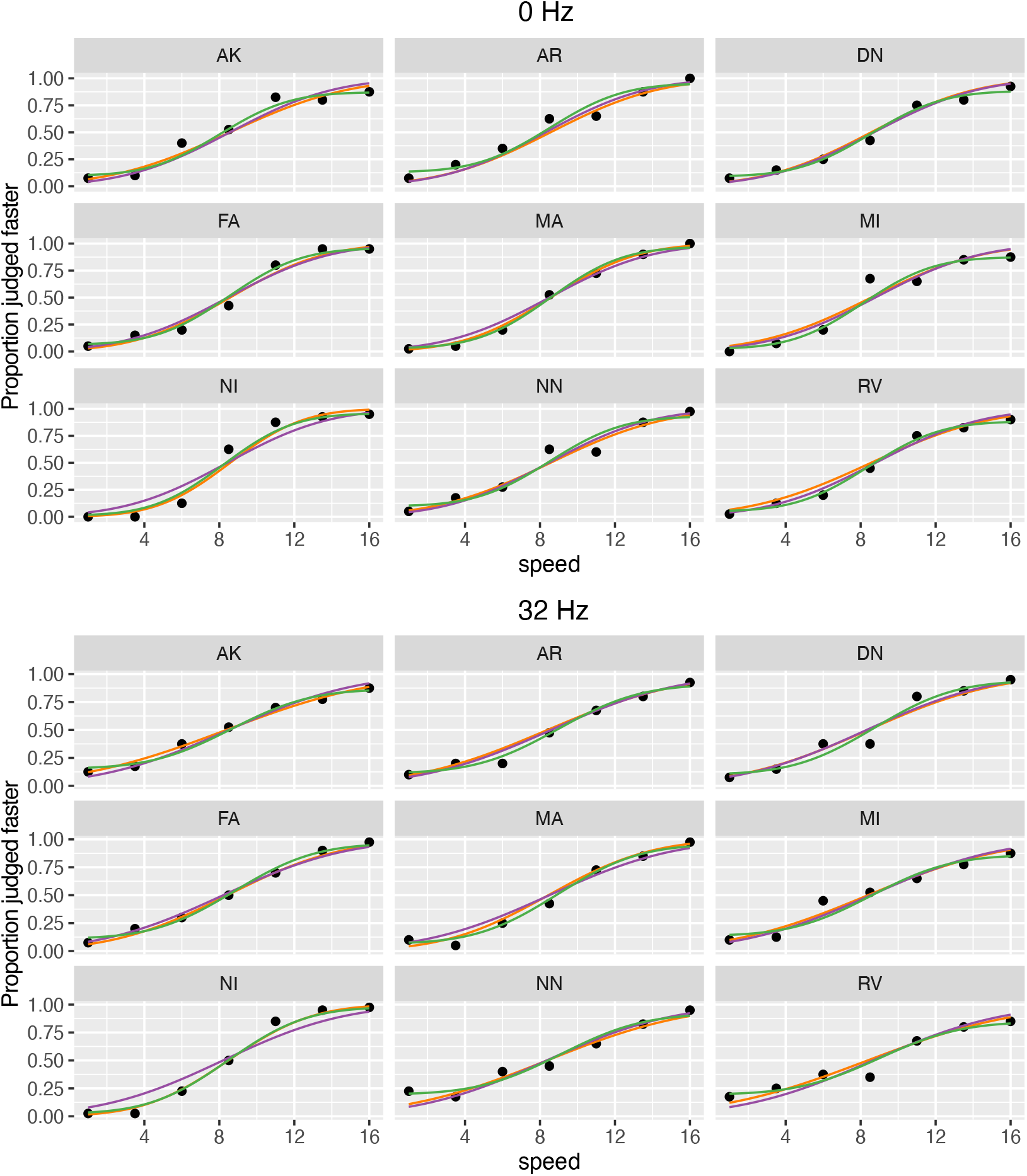
Hierarchical model predictions for speed discrimination data: GLMM (orange lines), BHF without lapse and guess rates (purple lines), and BHF with subject-specific lapse and guess rates (green lines). Proportions are shown as black points.

The predictors shown in Table 5 correspond to the coefficients *β*_0_, …*β*_3_ in Eq. (12). The first and second estimates represent the intercept and slope in the baseline condition (i.e. in the absence of masking vibration), while the third and fourth estimates capture the change in intercept and slope, respectively, associated with the presence of masking vibration (32 Hz condition). All fixed effects are statistically significant, with p-values from the *lmerTest* package falling below the threshold of *α* = 0.05. This suggests that the fixed effects included in the model meaningfully contribute to explaining the variability in the response.

It is possible to use functions from the MixedPsy package to estimate PSE and JND with the Delta Method:

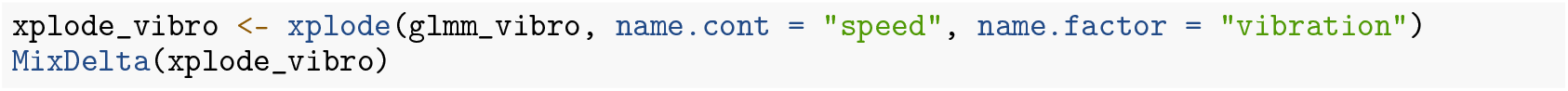

To compute bootstrap-based estimates of the parameters, it is necessary to account for the structure of the dummy-coded coefficients. In particular, the coefficients associated with the second level of the `vibration` condition (32 Hz) are obtained by summing the dummy-coded effect estimates with their respective baseline counterparts. This requires defining a function that calculates the PSE and JND from the fixed-effect estimates of the model as shown in Eqs. (13) and (14):

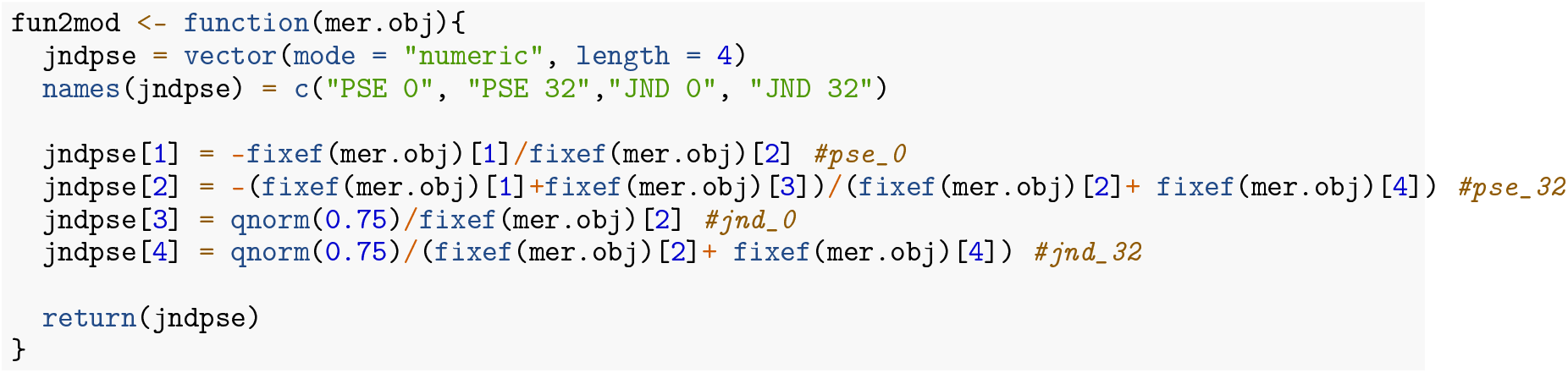

The custom function `fun2mod` must be provided as an input argument to the `pseMer()` function from the *MixedPsy* package in order to perform bootstrap-based estimation of the psychometric parameters:

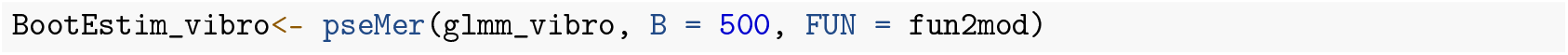

The values obtained from the bootstrap procedure are shown in Supplementary Table 1. The `fun2mod` function can be used to perform bootstrapping for other parameters if interest. For example, the following function extracts slopes for the two conditions (0 Hz and 32 Hz) and their difference (32 Hz – 0 Hz):

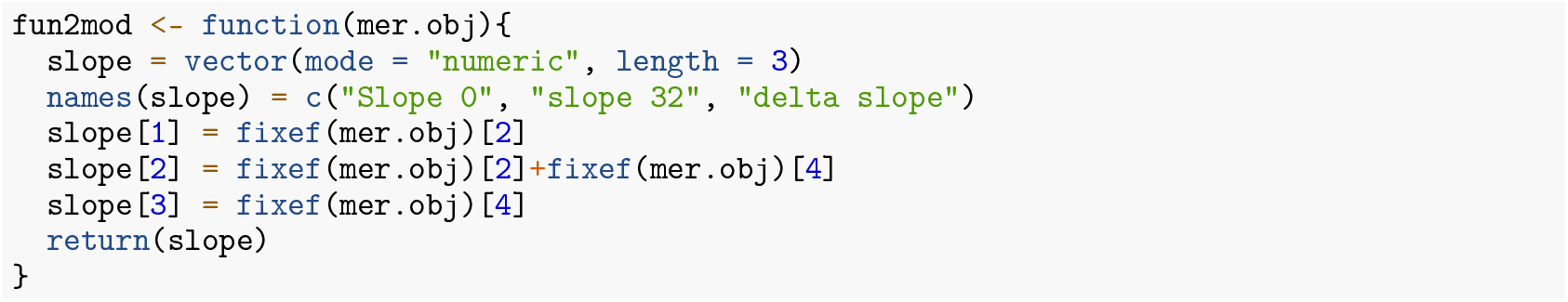

#### BHF

The Stan code used to fit the experimental dataset is largely similar to that presented in the previous sections for simulated data. Complete Stan programs (for models with and without lapse and guess rates), together with the R scripts for compiling them, are available for download. The main adjustments concern the `data` block, which must be modified to match the variable names of the `vibro_exp3` dataset, and the presence of the vibration condition with two levels, which require changes in the parametrization of the model. In particular, the `parameters` block must provide separate values for each vibration condition:

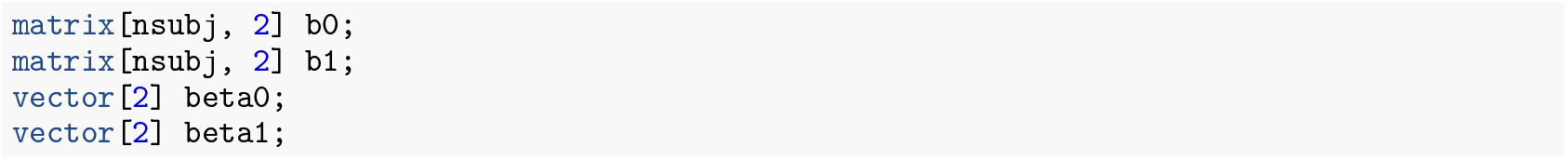

For this dataset, we adopted the parametrization in Eq. (15), focusing on the slope of the response as in the original paper (Dallmann et al., 2015). Accordingly, the `transformed parameters` block computes the linear predictor and the response probabilities as:

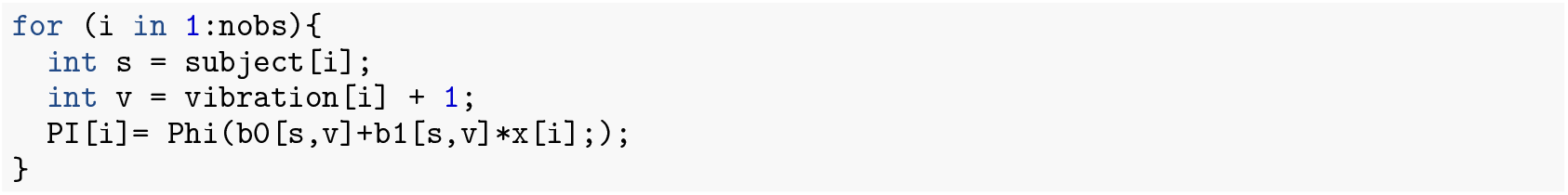

In addition, we define quantities that are useful for subsequent analysis, such as group-level and individual-level PSEs and JNDs, and the difference between slopes across vibration conditions:

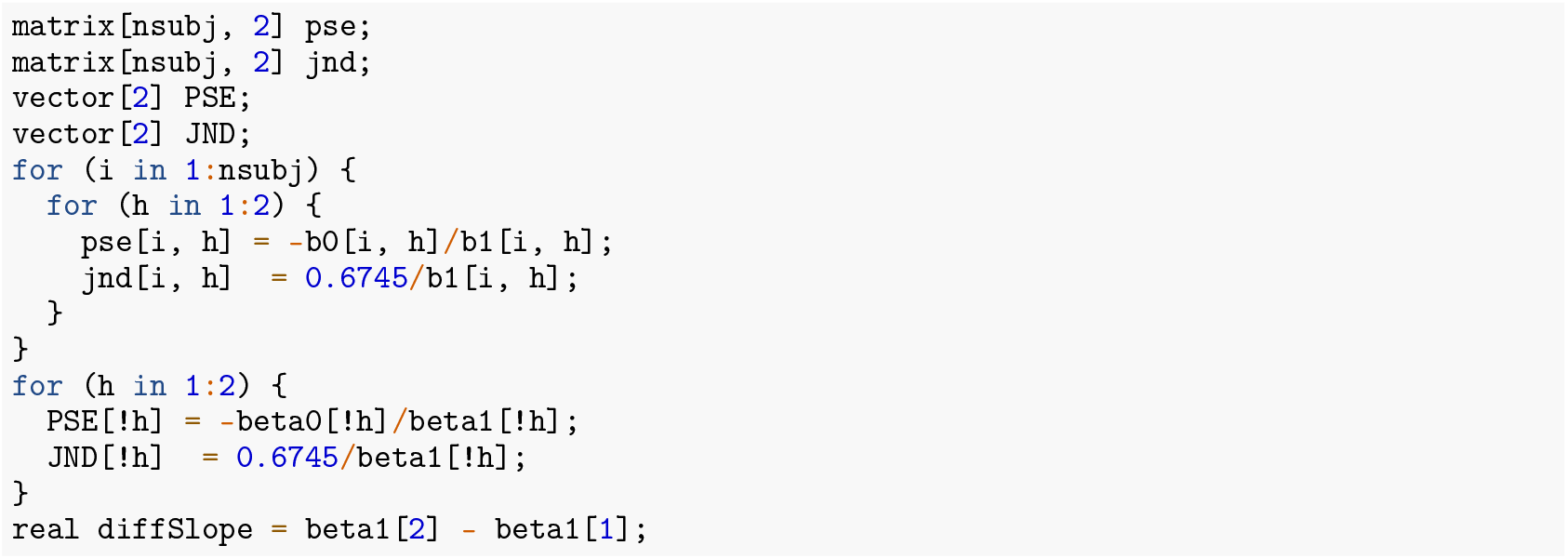

Finally, the `model` block accounts for the two vibration conditions by specifying condition-specific priors for the hierarchical parameters:

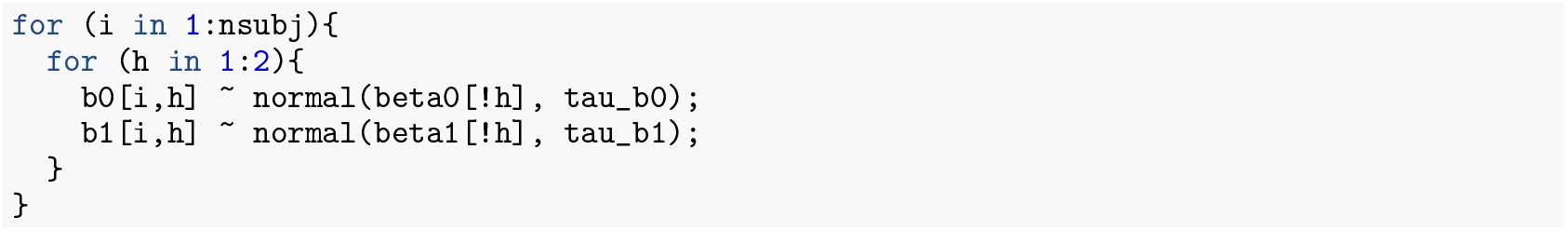

As shown in Example 1, the model can be extended to include individual guess and lapse rates for each vibration condition. This is achieved by introducing the corresponding parameters in the `parameters` block:

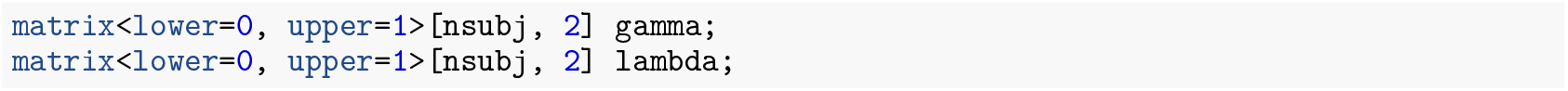

The response probability must then be modified accordingly, following the relations defined in Eq. 25. In particular, for each observation the probability becomes:

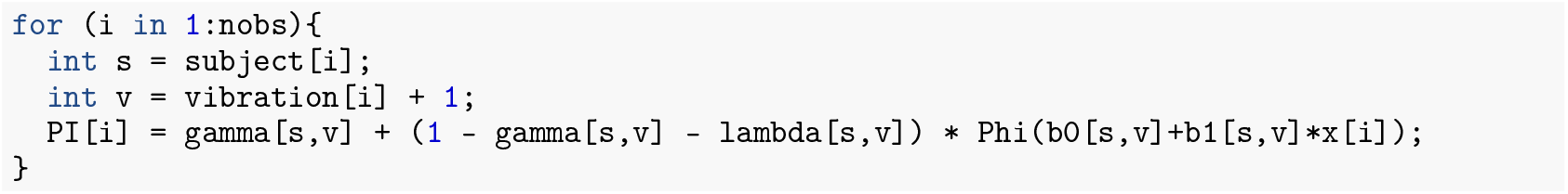

For this dataset, the initialization function defined for the call to `stan` accounts for the two levels of the vibration condition:

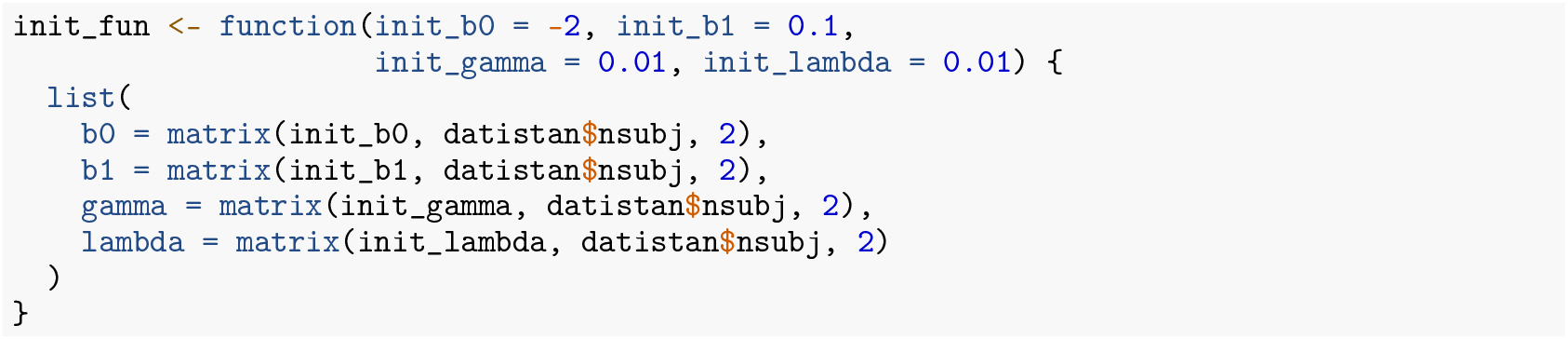

Model predictions for the two BHFs are shown Fig. 6 (purple lines for BH-GLM, green lines for BH-GNM).

### Model Diagnostics and Statistical Inference

Visual inspection of both the single-subject and hierarchical model predictions (Fig. 5–6) does not reveal substantial differences across models, aside from a few individual cases. The SSE values (Table 6) are also broadly comparable, with the BH-GNM showing slightly lower SSE–consistent with its additional parameters. DHARMa diagnostics (Supplementary Fig. 2) do not indicate major issues for any of the hierarchical models, aside from minor quantile deviations for the BH-GLM (bottom panel). Overall, the inclusion of guess and lapse parameters is not necessary in this dataset. Since no features requiring Bayesian modeling (e.g., power priors) are used, the GLMM provides an adequate choice in this case.

**Table 6.** SSE for residuals of hierarchical models in Example 2.

|  | Model | SSE |
| --- | --- | --- |
| 1 | GLMM | 0.421 |
| 2 | BH-GLM | 0.569 |
| 3 | BH-GNM | 0.361 |

For statistical inference, in line with the original study by Dallmann et al. (2015), we limited our analysis on the modulatory effect of masking vibration on the slope of the response. Estimates of PSE and JND are reported in Supplementary Table 1. We considered slope estimates across models (Fig. 7) and assessed the effects of vibration on slope differences (Fig. 8).

**Figure 7.**
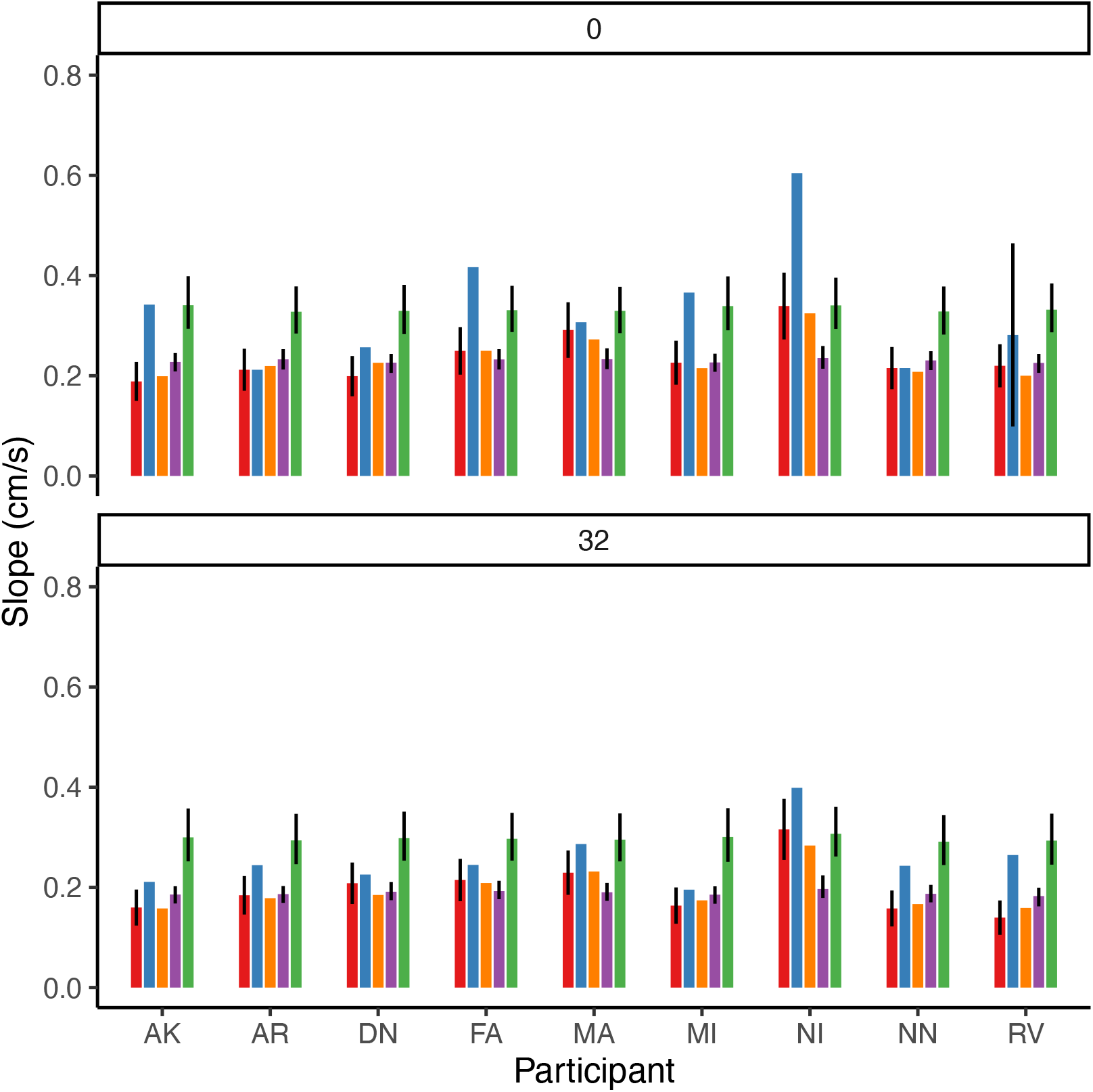
Slope estimates by participant and vibration condition. Colors indicate the model used: red = GLM, blue = GNM, orange = GLMM, purple = BH-GLM, green = BH-GNM. Error bars show 95% uncertainty intervals: confidence intervals based on Delta method for GLM, credible intervals for the BHF. GLMM does not provide a measure of uncertainty for individual participants. Bootstrap-based intervals for GNM are not shown, as they exceeded the figure boundaries due to lack of convergence of the bootstrap procedure.

**Figure 8.**
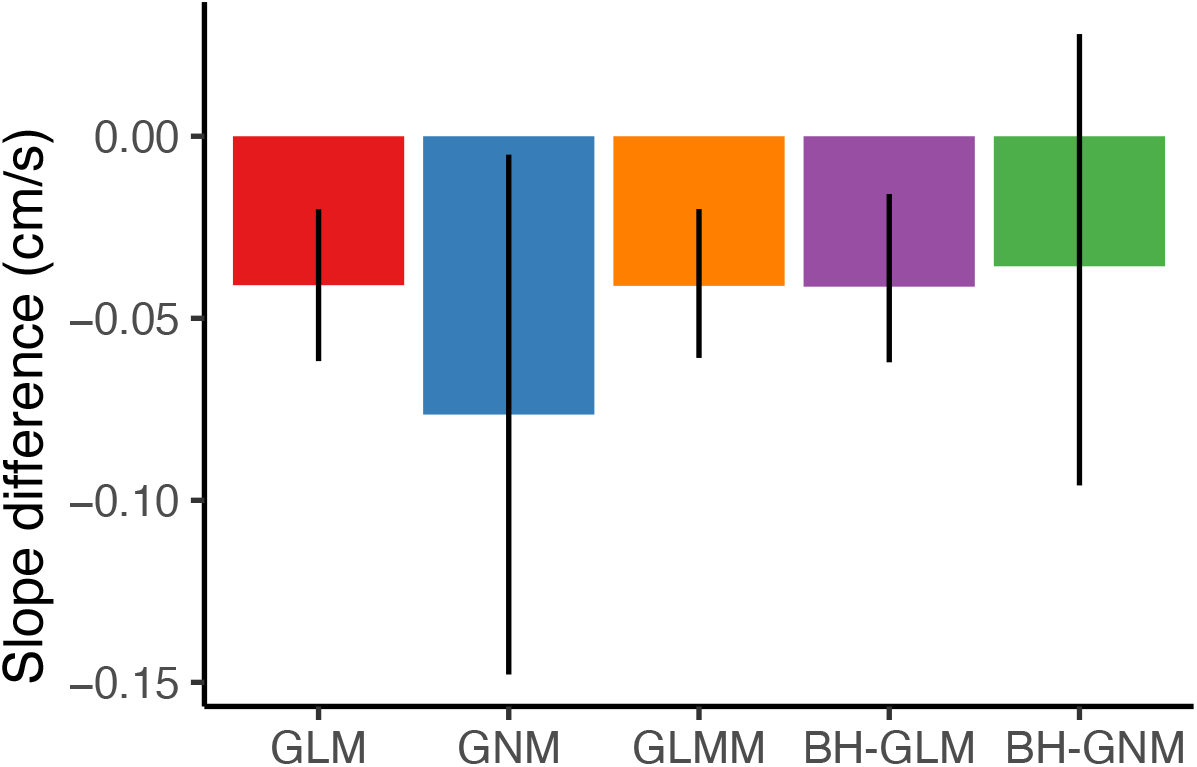
Slope differences (32 – 0 Hz) across the 5 models. Error bars represent 95% uncertainty intervals: sample-based confidence interval for GLM and GNM; bootstrap-based confidence interval (500 iterations) for GLMM; credible intervals for the BHFs.

For single-subject models (GLM and GNM) where a two-level approach was required, paired samples t-tests showed significant differences between the two conditions, with the slope at 32 Hz being smaller than at 0 Hz (GLM: mean difference = −0.041, t = −4.521, p = 0.002; GNM: mean difference = −0.076, t = −2.74, p = 0.039).

For the GLMM, the bootstrap-based CI of the slope difference lay below 0 (Estimate =−0.041, 95% CI = [−0.061, −0.020]), suggesting a significant effect of the vibration on the slope. Accordingly, the change in slope was a significant fixed effect on the model (Estimate *±* SE: −0.041 *±* 0.010, p < 0.001).

For Bayesian models, slope differences were computed as transformed parameters within the Stan program to obtain credible intervals. In the BHF without lapse and guess rates, the CrI for the slope difference lay entirely below 0 (Estimate = −0.041, 95% CrI = [−0.062, −0.016]). In contrast, when lapse and guess rates were modeled as random effects, the CrI was wider and included 0 (Estimate = −0.036, 95% CrI = [−0.096, 0.028]).

As in Example 1, *Shinystan* package confirms the stability of the sampling chains and, consequently, of the posterior estimates. 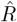 statistics always assumes values below 1.02 for the posterior distributions, after all the parameters in both the BH-GLM and BH-GNM models, after 15000 iterations with a burn-in of 10000. A negligible number of sampling warnings after warm-up were detected.

## Discussion

Several papers in the psychophysics literature adopt a two-level approach, in which a given parameter (e.g., *PSE*_*i*_) is first estimated separately for each participant *i*, and statistical inference is then performed on these individual estimates using a t-test or ANOVA, depending on the experimental design. However, this procedure is problematic because it ignores the standard errors associated with each individual estimate, leading to potentially biased or inefficient inferences. Hierarchical models provide a more appropriate alternative, as they correctly account for both within- and between-subject variability.

In this tutorial, we evaluated four model frameworks for fitting and analyzing psychophysical data: the Generalized Linear Model (GLM), the Generalized Nonlinear Model (GNM), the Generalized Linear Mixed Model (GLMM), and the Bayesian Hierarchical Model Framework (BHF) with two different parametrization (BH-GLM, BH-GNM). In the GLM and GNM frameworks, inference at population level is based on the two level approach detailed above. We compared their performance using two datasets from forced-choice discrimination tasks. The first dataset was simulated, with small but non-zero lapse and guess rates; the second was an experimental dataset from (Dallmann et al., 2015), in which we previously reported, using a GLMM, a significant difference in the parameter of slope between two experimental conditions. The experimental dataset was also analyzed in our previous article (Mezzetti et al., 2023).

In the simulated dataset, all models estimated the PSE accurately. However, models that did not account for lapse and guess rates (GLM, GLMM, BH-GLM) systematically overestimated the JND compared with models that incorporated these parameters (GNM, BH-GNM). The latter yielded JND estimates very close to the simulated values. The underestimation of the JND when non-zero lapses and guesses are not accounted by the model is in accordance with Wichmann & Hill (2001a). In the experimental dataset, all models produced comparable fits, and parameter estimates were consistent across frameworks (see Figures 7–8 and Supplementary Table 1). All models except the BH-GNM detected the previously reported difference in slope between conditions. When the effect of lapsing and guessing is negligible, the models with less parameters are usually preferred.

Each of the evaluated models has distinct advantages and limitations. Understanding these differences is essential for selecting the most appropriate approach for a given analysis. In Fig. 9 we proposed a possible flowchart to guide the researchers in the choice of the most appropriate model.

**Figure 9.**
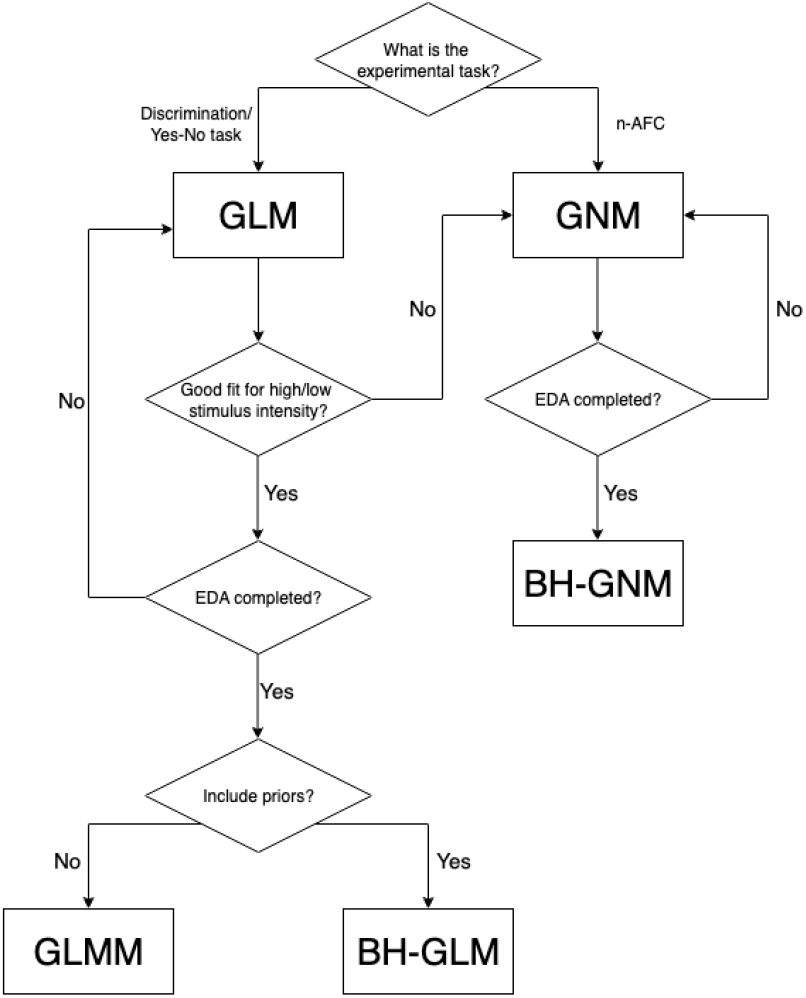
Example of analysis pipeline with the proposed model frameworks.

In GNM and BH-GNM we approximated the equation of the PSE and JND without taking into account the correction term depending on *λ* and *γ*. The correction term is negligible when *λ ≈ γ*; for example assuming a difference between the two equal to 0.003, the correction to the PSE is 0.004. In typical n-AFC experiments, the threshold stimulus *x*_*ij*_(*p*) corresponds to the 50th percentile of the psychometric function. For example, in a 2-AFC task, where the lower asymptote *γ* = 0.5, assuming *λ* = 0 the threshold is *x*_*ij*_(0.75). Importantly, for the 50th-percentile threshold, we always have *x*_*ij*_(*p*) = *µ*_*i*_.

Models fitted at the single-subject level (GLM, GNM) are straightforward to implement and computationally light. The GNM framework, in particular, offers flexibility for customizing model structure and adding coefficients as needed. However, as previously demonstrated by Moscatelli et al. (2012), two-level approaches (individual fits followed by group-level inference) do not retain within-subject variability in second-level analyses and may have lower statistical power compared to hierarchical models. Moreover, GNMs can suffer from convergence difficulties and unreliable standard error estimation due to their non-linear structure and reliance on maximum likelihood methods. With limited data, bootstrapped standard errors may also be unstable because of these convergence issues.

Frequentist mixed models such as GLMMs overcome some of these limitations by increasing statistical power and accounting for both between- and within-subject variability. However, a key limitation of GLMMs is their inability to estimate lapse and guess rates. This lack of control over response asymptotes can be problematic, particularly in n-AFC tasks—where the lower asymptote is fixed at 1*/n* or when data exhibit high lapse rates (Wichmann & Hill, 2001a). Although it is possible to model the variability between participants by means of random-effect parameters, GLMMs do not provide estimates of the variance within individual participants (as illustrated in Fig. 3 and 7). It is important to note that GLMMs do not rely on the statistical assumptions of asymptotic normality of the parameters (Agresti, 2002; Casella & Berger, 2002). For this reason, simulations are generally recommended over analytic methods for conducting inferential statistics on parameters of interest (James et al., 2013; Moscatelli et al., 2012). Given the higher computational demands of bootstrap estimation, the delta method should be used only for preliminary evaluation of effects and subsequently confirmed using the bootstrap approach when reporting final results.

The Bayesian Hierarchical Framework (BHF) offer the greatest flexibility, allowing full specification of model structure, parameter constraints, and prior distributions, making them the most suitable option when modeling guess and lapse rates. In the BHF, random effects are treated as estimated parameters, and credible intervals can be computed to quantify uncertainty both for individual-level estimates and for population-level (fixed) effects. As shown in our previous work (Mezzetti et al., 2023), the BHF can incorporate informative or power priors to integrate prior knowledge from earlier studies. This flexibility comes at the cost of greater computational demands and the need for careful model specification. An important consideration when choosing a modeling approach is that Bayesian inference differs fundamentally from frequentist inference. In the Bayesian framework, all parameters are treated as random variables with their own probability distribution (Zhao et al., 2006; Fong et al., 2010). Instead of producing a point estimate with associated p-value, Bayesian models yield a posterior distribution for each parameter, allowing a direct and natural assessment of uncertainty. Consequently, traditional frequentist metrics such as p-values are not directly available; inference is expressed in terms of posterior probabilities, credible intervals, or other probabilistic summaries of the parameters. This approach can be unusual and more difficult to interpret for researchers who are accustomed to working with p-values and null-hypothesis significance testing, requiring a shift in statistical thinking and reporting conventions.

Model selection should be guided by the characteristics of the data and the specific goals and constraints of the analysis. For Exploratory Data Analysis (EDA) or small datasets in pilot studies, single-subject models (GLM or GNM) are often sufficient—with GNM being the most appropriate solution when behavior at the asymptotes of the curve warrants explicit modeling of lapse and/or guess rates (Fig. 9). When data are ample and hierarchical modeling of variability is desired, GLMMs represent an excellent compromise, offering robust inference and interpretability within a frequentist framework. Notably, as shown in the proposed examples, GLMMs and BH-GLMs with non-informative priors produce nearly identical estimations. Bayesian models are most appropriate when prior information is available (as in the examples described in our previous work (Mezzetti et al., 2023)), model customization is needed, or full probabilistic inference is desired. Ultimately, the aim is to achieve an appropriate balance between model complexity and parsimony—avoiding overfitting while adequately capturing the key features of the data James et al. (2013).

This tutorial provided general guidelines and recommendations, exemplified by the workflow shown in Fig. 9. However, the selection of the most appropriate model ultimately depends on the specific objectives of the study and the characteristics of the dataset, and thus requires the researcher’s informed and context-sensitive judgment.

## Acknowledgments

We would like to thank Francesco Lacquaniti for useful comments and suggestions on the manuscript. This study was partially supported by the Italian Ministry of Health (Ricerca corrente, IRCCS Fondazione Santa Lucia) and the European Commission H2020 Framework Program (HARIA, Grant No. 101070292).

## Data Availability

The data and materials for the tutorial are available for download on CRAN and GitHub. None of the experiment was preregistered.

