## Supplemental materials for "Modeling Psychophysical Data in R: A Comparative Study of Four Model Frameworks"

### Supplementary Material

A Preprint

**Priscilla Balestrucci**
Laboratory of Neuromotor Physiology
Santa Lucia Foundation IRCCS, Rome, Italy


**Maura Mezzetti**
Department of Economics and Finance
University of Rome Tor Vergata, Rome, Italy


**Barbara La Scaleia**
Laboratory of Neuromotor Physiology
Santa Lucia Foundation IRCCS, Rome, Italy


**Alessandro Moscatelli** *
Department of Systems Medicine and Centre of Space Biomedicine
University of Rome Tor Vergata, Rome, Italy
Laboratory of Neuromotor Physiology
Santa Lucia Foundation IRCCS, Rome, Italy


November 26, 2025

#### Overview

This document contains supplementary information supporting the manuscript entitled:

*Modeling Psychophysical Data in R: A Comparative Study of Four Model Frameworks.*

*Corresponding author

### Additional Figures

#### DHARMa residuals - Example 1

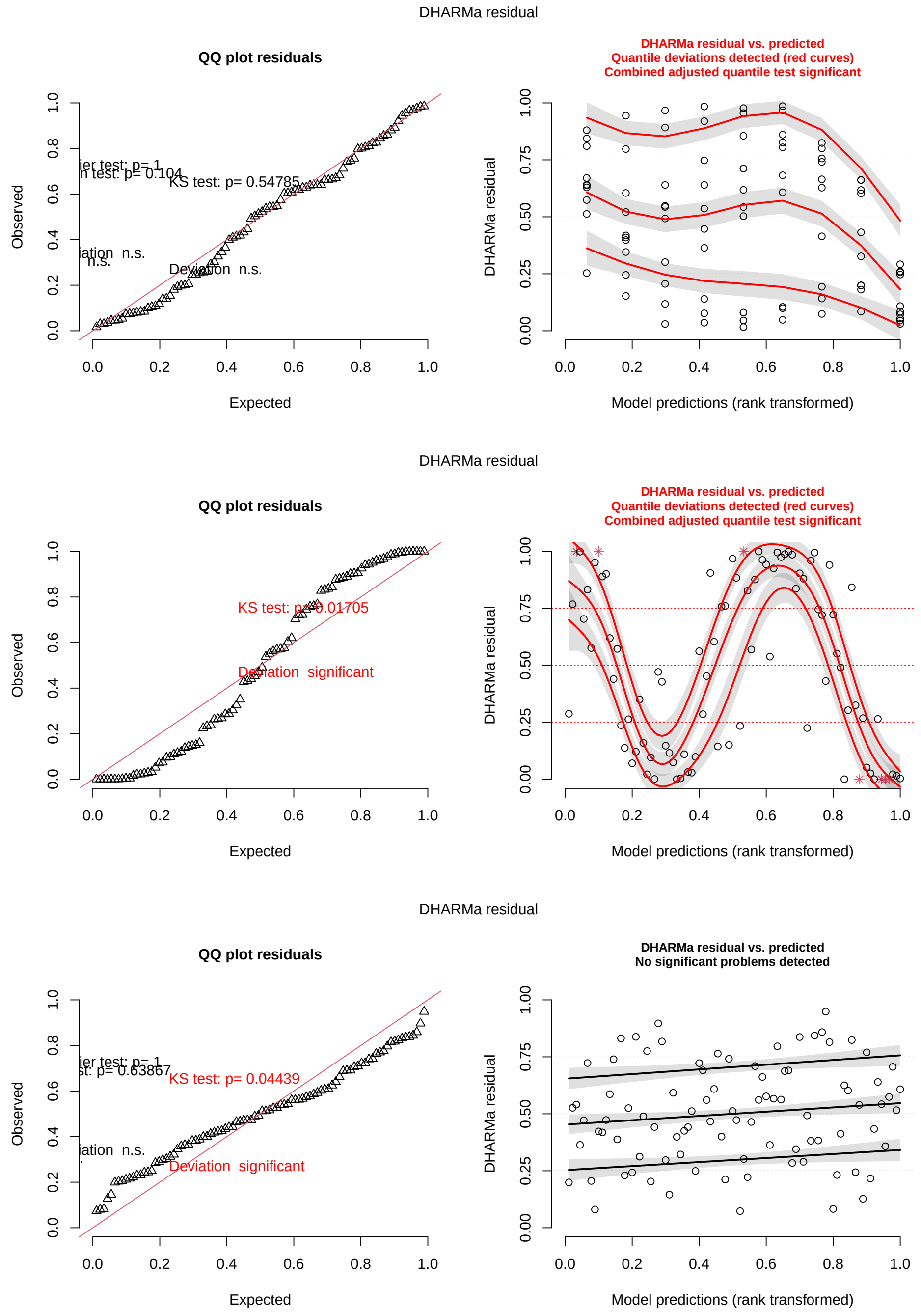


Figure 1: DHARMa residuals for hierarchical model fitted to a simulated dataset. Top: GLMM; Middle: BH-GLM; Bottom: BH-GNM. Refer to Table 1 for acronyms.

#### DHARMa residuals - Example 2

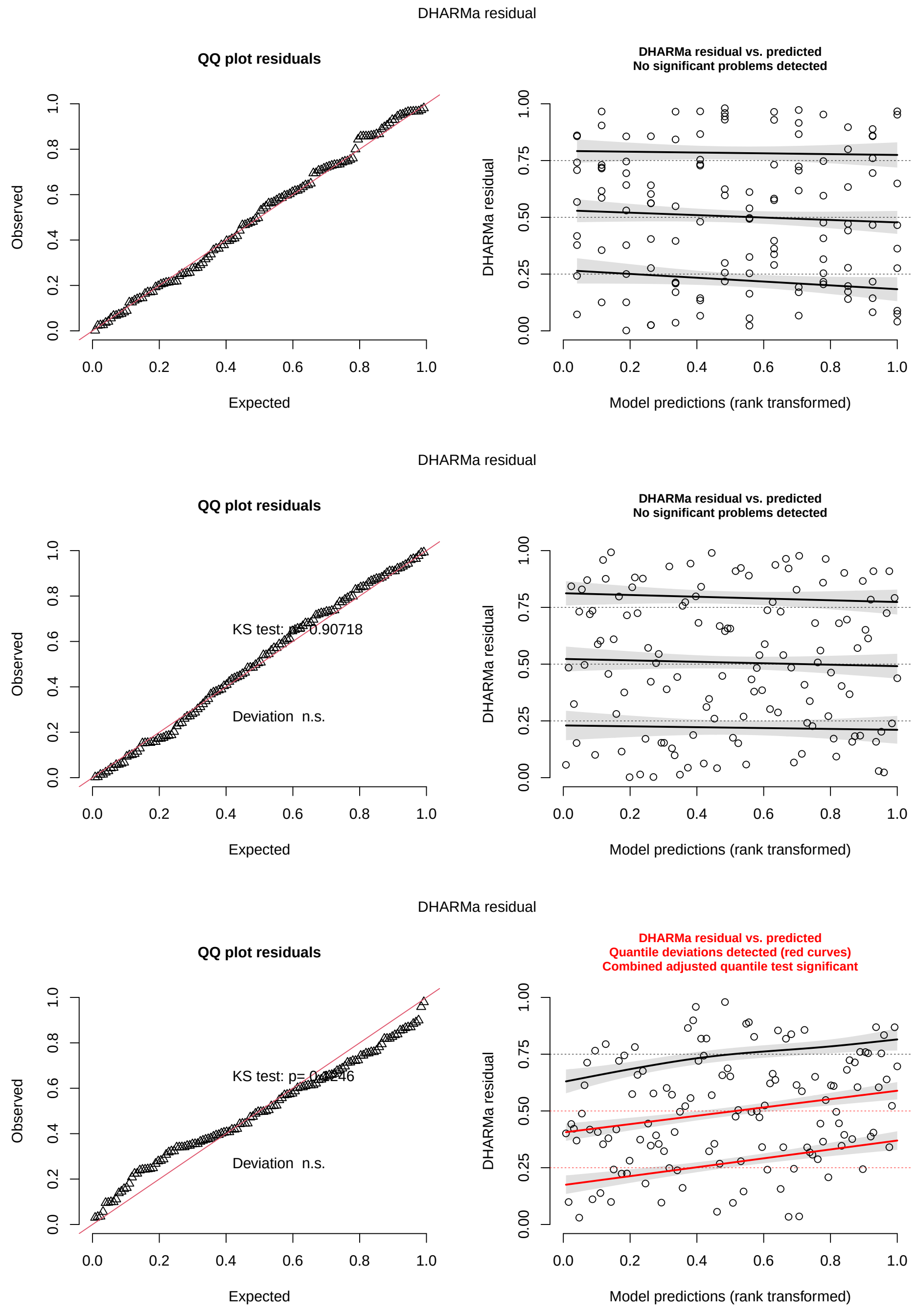


Figure 2: DHARMa residuals for hierarchical model fitted to dataset (vibro). Top: GLMM; Middle: BH-GLM; Bottom: BH-GNM.Refer to Table 1 for acronyms.

#### PSE and JND estimates for data in Example 2

Table 1: Mean and 95% uncertainty intervals (top: PSE, bottom: JND) of psychometric parameters for the data in Example 3, estimated by the five models. Uncertainty intervals are sample-based intervals for GLM and GNM; bootstrap-based confidence intervals (500 iterations) for GLMM; credible intervals for the BH-GLM and BH-GNM. Refer to Table 1 for acronyms.

| models | vibration 0 | vibration 32 |
|---|---|---|
| GLM | 8.565 [7.711, 9.419] | 8.409 [7.821, 8.997] |
| GNM | 8.184 [6.890, 9.478] | 8.862 [7.073, 10.650] |
| GLMM | 8.565 [8.329, 8.847] | 8.416 [8.120, 8.746] |
| BH-GLM | 8.550 [8.171, 8.932] | 8.423 [7.956, 8.867] |
| BH-GNM | 8.379 [7.880, 8.868] | 8.795 [8.232, 9.430] |

| models | vibration 0 | vibration 32 |
|---|---|---|
| GLM | 2.926 [1.930, 3.921] | 3.619 [1.985, 5.252] |
| GNM | 2.231 [0.892, 3.571] | 2.725 [1.730, 3.721] |
| GLMM | 2.864 [2.511, 3.283] | 3.469 [2.982, 4.089] |
| BH-GLM | 2.936 [2.753, 3.174] | 3.579 [3.325, 3.846] |
| BH-GNM | 2.034 [1.777, 2.311] | 2.283 [1.948, 2.643] |

#### PSE and JND in GNM model types

The PSE is the stimulus value yielding to a probability $\pi(x) = 0.5$. Substituting in Eq. 8 gives

$$0.5 = \gamma_i + (1 - \gamma_i - \lambda_i)F(x_{ij}; \mu_i, \sigma_i)$$

Solving for $x_{ij} = PSE_i$:

$$\begin{aligned} 0.5 &= \gamma_i + (1 - \gamma_i - \lambda_i)F(PSE_i; \mu_i, \sigma_i) \\ 0.5 - \gamma_i &= (1 - \gamma_i - \lambda_i)F(PSE_i; \mu_i, \sigma_i) \\ \frac{0.5 - \gamma_i}{1 - \gamma_i - \lambda_i} &= F\left(\frac{PSE_i - \mu_i}{\sigma_i}\right) \end{aligned}$$

which leads to:

$$PSE_i = \mu_i + F^{-1}\left(\frac{0.5 - \gamma_i}{1 - \gamma_i - \lambda_i}\right)\sigma_i$$

We can define:

$$k_i(\gamma_i, \lambda_i) = F^{-1}\left(\frac{1}{2}\frac{1 - 2\gamma_i}{1 - \gamma_i - \lambda_i}\right)\sigma_i$$

so that

$$PSE_i = \mu_i + k_i$$

If $\gamma_i \approx \lambda_i$, as is typical in forced-choice discrimination experiments, the shift $k_i$ is small, therefore $PSE_i \approx \mu_i$. If $\gamma_i = \lambda_i$, $k_i = 0$ and $PSE_i = \mu_i$.

The PSE is defined as the stimulus value corresponding to a response probability $p = 0.5$ and is used in forced-choice experiments where the response probability spans approximately 0 to 1.

Instead, in n-AFC tasks, the lower asymptote is $1/n$, and the threshold $x_i$ can be calculated corresponding to the stimulus value yielding to the 50th percentile of the psychometric function. For example, in a 2-AFC task where $\gamma = 1/2$ and $\lambda = 0$, the threshold $x_i$ is the stimulus values that corresponds to $\pi(x_i) = 0.75$. Substituting into the equations above, it can be seen that in this case $k_i = 0$, so that $x_i = \mu_i$.

For the JND, we use the definition in Eq. 4. Given $\sigma_i = 1/\beta_{1i}$, we have:

$$JND_i(p) = \sigma_i \left[ F^{-1}\left(\frac{p - \gamma_i}{1 - \gamma_i - \lambda_i}\right) - F^{-1}\left(\frac{0.5 - \gamma_i}{1 - \gamma_i - \lambda_i}\right) \right]$$

Using the definition of $k_i$, this can be rewritten as

$$JND_i(p) = \sigma_i F^{-1}\left(\frac{p - \gamma_i}{1 - \gamma_i - \lambda_i}\right) - k_i$$

For small values of $\gamma_i, \lambda_i$, the JND can be approximated by

$$JND_i(p) \approx \sigma_i F^{-1}(p)$$
